# ASAP: An automatic sustained attention prediction method for infants and toddlers using wearable device signals

**DOI:** 10.1101/2024.11.16.623211

**Authors:** Yisi Zhang, A. Priscilla Martinez-Cedillo, Harry T. Mason, Quoc C. Vuong, M. Carmen Garcia-de-Soria, David Mullineaux, Marina Knight, Elena Geangu

**Affiliations:** Department of Psychological and Cognitive Sciences, Tsinghua University, Beijing, 100084, People’s Republic of China; Tsinghua Laboratory of Brain and Intelligence (THBI), Tsinghua University, Beijing, 100084, People’s Republic of China; Department of Psychology, University of York, York, England YO10 5DD; Department of Psychology, University of Essex, Wivenhoe Park, Colchester, Essex, England CO4 3SQ; School of Physics, Engineering and Technology, University of York, York, England YO10 5DD; Bioscience Institute, Newcastle University, Newcastle Upon Tyne, England NE1 7RU; School of Psychology, Newcastle University, Newcastle Upon Tyne, England NE1 7RU; Department of Mathematics, University of York, York, England YO10 5DD

## Abstract

Sustained attention (SA) is a critical cognitive ability that emerges in infancy. The recent development of wearable technology for infants enables the collection of large-scale multimodal data in the natural environment, including physiological signals. To capitalize on these new technologies, psychologists need methods to efficiently extract valid and robust SA measures from large datasets. In this study, we present an innovative automatic sustained attention prediction (ASAP) method that harnesses electrocardiogram (ECG) and accelerometer (Acc) signals recorded with wearable sensors from 75 infants (6-, 9-, 12-, 24- and 36-months). Infants undertook various naturalistic tasks similar to those encountered in their natural environment, including free play with their caregivers. Annotated SA was validated by fixation signals from eye-tracking. ASAP was trained on temporal and spectral features derived from the ECG and Acc signals to detect attention periods, and tested against human-coded SA. ASAP’s performance is similar across all age groups, demonstrating its suitability for studying development. We also investigated the relationship between attention periods and low-level perceptual features (visual saliency, visual clutter) extracted from the egocentric videos recorded during caregiver-infant free play. Saliency increased during attention vs inattention periods and decreased with age for attention (but not inattention) periods. Crucially, there was no observable difference in results from ASAP attention detection relative to the human-coded attention. Our results demonstrate that ASAP is a powerful tool for detecting infant SA elicited in natural environments. Alongside the available wearable sensors, ASAP provides unprecedented opportunities for studying infant development in the ‘wild’.

## Introduction

Sustained attention (SA) is a fundamental cognitive ability that emerges during infancy and has a widespread impact on various developmental domains (Reynolds & Romano, 2016; Rose et al., 2001; Ruff & Lawson, 1990). As a form of endogenous attention, it refers to the ability to focus on a particular stimulus or event over an extended period even with distractors present (Casey & Richards, 1988). SA shapes infant learning (Brandes-Aitken et al., 2023; Frick & Richards, 2010; Xie et al., 2017) and is associated with the emergence in infancy and early childhood of more complex cognitive and social abilities, such as executive function (Brandes-Aitken et al., 2023; Johansson et al., 2015), self-regulation (Brandes-Aitken et al., 2019; Casey & Richards, 1988; Rothbart et al., 2011; Ruff, 1986; Swingler et al., 2015), memory (Nelson & deRegnier, 1992; Reynolds & Romano, 2016; Richards, 2003), language, and social communication (Mundy & Jarrold, 2010; Salley et al., 2016).

Most developmental SA research relies on lab-based paradigms that fail to capture the complexity of infants’ day-to-day environment (Bradshaw et al., 2024; Brandes-Aitken et al., 2019; Xie et al., 2018), significantly limiting the ecological validity of any proposed theories (Dahl, 2016). The development of affordable, lightweight and user-friendly wearable technology for infants has facilitated the recording of what infants see and hear in natural settings, and related autonomic nervous system (ANS) response (Geangu et al., 2023). These have great potential for precise measurement and characterisation of SA development as a function of mutual interactions with various factors (Zhu et al., 2015). However, a key challenge in adopting ecologically valid approaches is the lack of efficient methods for extracting reliable measures of SA from the resulting vast amount of data (Long et al., 2022). In this study, we reduce this bottleneck by introducing an innovative automatic attention detection algorithm that harnesses infant electrocardiography (ECG) and accelerometer (Acc) signals recorded with wearable sensors.

Traditionally, looking times have been the predominant measure for visual attention in infancy (Aneja et al., 2024; Kadooka & Franchak, 2020; Papageorgiou et al., 2014). However, brain imaging and psychophysiology evidence indicate that looking times cannot accurately differentiate infant visual SA from other attentional processes and inattention (Colombo, 1995; Richards, 2003; Richards & Turner, 2001). Characterisation of heart rate (HR) deceleration and acceleration patterns provides a more reliable and accurate measure of attentional processes, particularly for visual SA (Richards, 2003; Richards & Casey, 1991; Xie et al., 2018). Orienting towards a stimulus of interest is typically characterized by rapid HR deceleration relative to baseline or prestimulus levels. If attention orientation is followed by SA for further processing, HR stabilises at the lower rate for 2-20 seconds, but can be even longer (Richards & Casey, 1991). This reduced HR is often also associated with a reduction in body movement (Friedman et al., 2005). Attention termination (disengagement from processing the stimulus of interest) is marked by a rapid HR acceleration to approximate pre-attention baseline levels (Brandes-Aitken et al., 2023). This infant HR-defined SA (HRDSA) is associated with frontal electroencephalogram (EEG) synchronisation in theta oscillations, which has been linked with goal-oriented attention in infancy (Brandes-Aitken et al., 2023; Xie et al., 2018), and has been observed for infants as young as 3 months (Brandes-Aitken et al., 2023). This supports HRDSA as a robust approach to investigating SA development.

Motivated by HRDSA, we establish an automatic SA prediction (ASAP) method to detect attention periods using ECG and Acc signals provided by wearable sensors used in infant research (Geangu et al., 2023; Islam et al., 2024; Maitha et al., 2020). This approach circumvents the need to use eye movements as predictors, acknowledging the challenges of capturing fixation data in natural environments. In contrast, the lab setting allows for accurate fixation measurements, enabling human coding and validation of HRDSA. Therefore, we generated a ground-truth dataset consisting of ECG and Acc signals, human-coded SA, and eye-tracking data in the lab. Our ASAP method is trained and validated using this dataset and can be applied to wearable sensor data collected in natural environments to study SA in a less intrusive manner.

ASAP can be decomposed into three primary steps, motivated by the abrupt HR deceleration and acceleration that occur during attention orientation and termination, immediately before and after the period of SA. The first step involves change point detection (CPD) (Aminikhanghahi & Cook, 2017; Fryzlewicz, 2020), which identifies time indexes indicating where the statistical properties of a signal undergo significant changes. Infancy and toddlerhood are characterised by frequent spans of SA (Slone et al., 2018; Yu & Smith, 2016), suiting a CPD algorithm which can objectively identify multiple change points within HR time series data. Secondly, momentary attention detection is formulated as a binary classification task. The classification model is trained with a curated set of features using a feature selection process. In the final step, the segmentation of SA periods is refined to preserve their temporal structure.

The temporal and spectral features of HR fluctuations provide rich information about the ANS’ regulation of homeostasis, physiological arousal, and cognitive states (Forte et al., 2019; Thayer et al., 2009). Precisely, heart rate variability (HRV)—measured through variations in beat-to-beat intervals in the time domain or through low (0.04-0.15 Hz) and high-frequency (0.15-0.4 Hz) HR oscillations in the frequency domain— is associated with sympathetic and parasympathetic nervous activity (Akselrod et al., 1981). Higher HRV is linked to better performance in SA tasks (e.g., continuous performance tests involving executive functions (Hansen et al., 2003)). Low-frequency HR oscillations are related to attentional demands (Börger et al., 1999; Van Roon et al., 2004), and an increased low-to-high frequency power ratio is associated with poor attention in children with ADHD (Griffiths et al., 2017). We characterise HR fluctuations in both time and frequency domains using wavelet transform to capture dynamic HRV changes, potentially relating to momentary shifts in attentional states. Wavelet analysis has been widely applied to characterise instantaneous features of physiological signals (Addison, 2005; Pichot et al., 1999; Toledo et al., 2003). Given the tight coordination between movement and cardiac output (Zhang et al., 2022), we also include Acc signals and investigate their dynamic coupling with HR in time and frequency for additional attention insights.

To demonstrate ASAP’s utility in attention research, we apply ASAP in a study examining what drives SA when infants and toddlers actively engage with people and objects. Attention allocation can be influenced by factors such as low-level visual features (e.g., strong edges, bright colours, large movements in the scene), people (e.g., faces and bodies), and objects (e.g., toys) (Amso et al., 2014; Byrge et al., 2014; Crespo-Llado et al., 2018; DeBolt et al., 2023; Geangu & Vuong, 2020, 2023; Kadooka & Franchak, 2020; Kwon et al., 2016; Oakes et al., 2024; Sun et al., 2024; Tummeltshammer et al., 2014; van Renswoude et al., 2019). However, most previous studies present images or videos under constrained lab conditions (e.g., sitting on a caregiver’s lap with stimuli presented on a monitor), which fail to capture infants’ and toddlers’ active interactions with their environment, and how this may change throughout development. Here we focus on visual saliency (Harel et al., 2006) and visual clutter (Rosenholtz et al., 2007) extracted from observers’ egocentric views during free play. Finally, we examine how these measures differ across fixated regions during attention and inattention periods, as determined by the ASAP method or human coders, and assess whether they vary with age.

## Methods

### Overview of approach

Our approach to developing an automatic SA-detection method hinges on two pivotal ideas: first, creating a lab-based ground-truth dataset with human-coded SA; second, training a model using this dataset to automatically detect SA based on ECG and Acc signals.

The multimodal dataset comprises four synchronised sensor signals: ECG, Acc, the scene videos, and fixation-label data. ECG and Acc signals are captured simultaneously from a wearable device attached to the infant’s chest. Fixation data are obtained from a head-mounted eye tracker that records infants’ and toddlers’ gaze points (Figure 1), which determine the objects fixated upon during recording. HR is derived from the ECG signal (Geangu et al., 2023; Mason et al., 2024). We annotated attention periods synchronised with the HR and Acc signals by considering both the distinctive HR waveforms during periods of SA and human inspection of aligned fixation data. (Figure 1, “Human Coding”).

**Figure 1.**
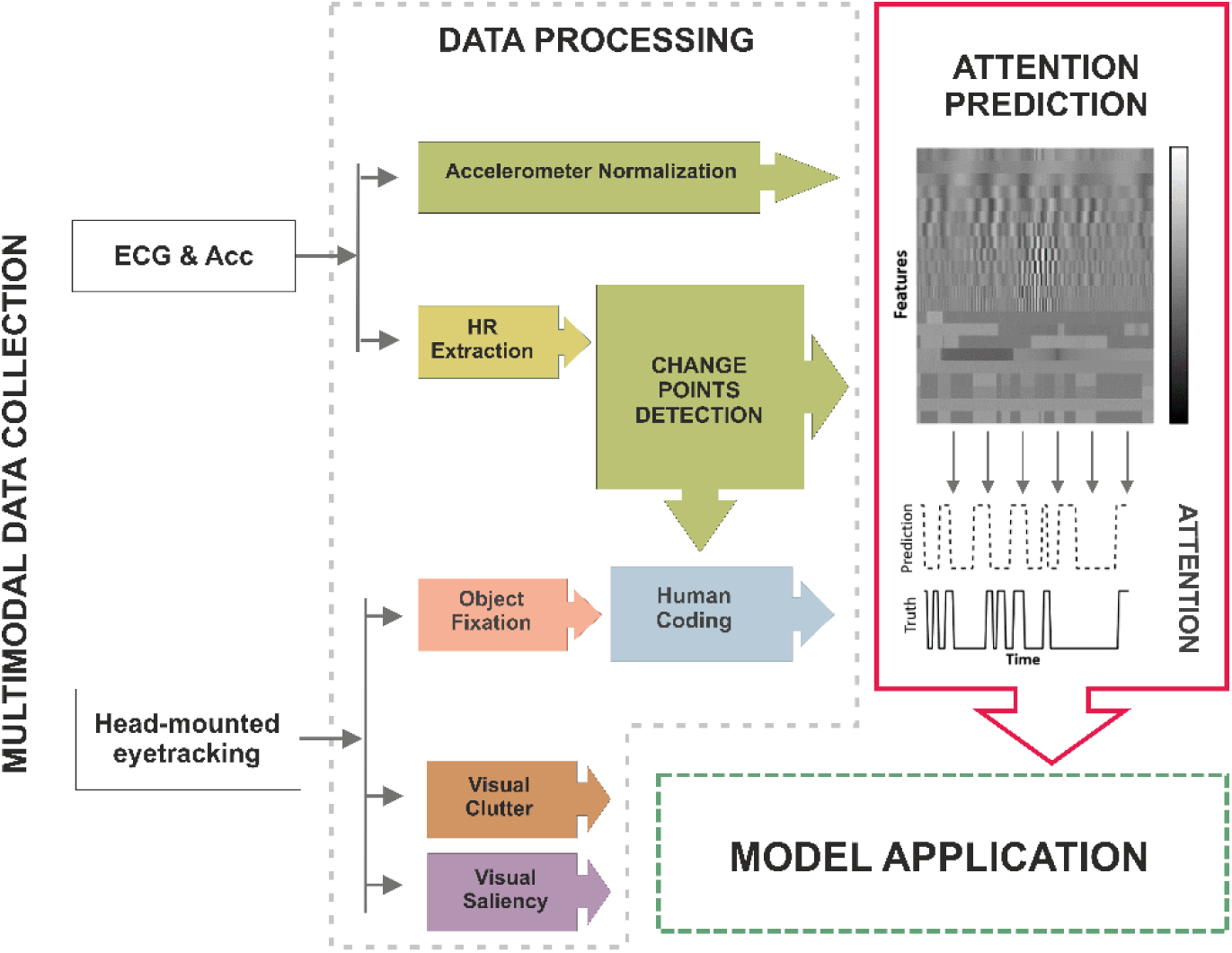
A pipeline schematic for data collection to train the attention prediction model. Two synchronised wearable devices record the data: the ECG/Acc body sensors and the head-mounted eye tracker. During data processing, HR is extracted and undergoes change points detection (CPD) to facilitate human coding of attention, validated by object fixation (objects coded by colours). During attention prediction, HR- and Acc-derived time series and change point segmentation form the feature matrix for training a machine learning model to predict attention periods. During model application, visual clutter and saliency signals are extracted from video frames to test the model’s effectiveness.

The model development uses HR and Acc signals alongside human-coded attention (Figure 1, “Attention Prediction”) and it involves three key stages: (1) coarse segmentation of potential periods of SA based on HRDSA facilitated by CPD; (2) classification employing HR and Acc features; and (3) fragment refinement to reconstruct the temporal structure. The model’s innovation lies in several aspects: first, the integration of a CPD algorithm to objectively identify abrupt changes in HR, adapting previous lab-based procedures (Richards & Casey, 1991) to naturalistic settings; second, the exploitation of temporal and spectral properties of HR and Acc signals and their interaction to delineate attentional states; and finally, preserving the temporal structure to retain the associated statistical characteristics throughout the attention detection process. We then demonstrate the model’s application in studying visual SA development by leveraging the egocentric video and fixation data collected with the HR and Acc signals (Figure 1, “Model Application”).

### The Dataset

#### Participants

Based on previous literature, N=75, 6- to 36-month-old infants and toddlers were included in the final analyses (Table 1). A further 23 participants responded to the invitation to participate in the study, but were not included in the final analysis due to refusal to wear one or all of the devices (*N* = 15) or technical errors (*N* = 8). Participants were recruited from an urban area in England. Caregivers provided written informed consent before the experimental procedure began, and families received £10 and a book. The research presented in this empirical report received ethical approval from the Department of Psychology Ethics Committee, University of York (Identification Number – 119). The experimental procedures were conducted in adherence to the principles of the1964 Declaration of Helsinki and its later amendments.

**Table 1:** The cohort breakdown of the participants in the Dataset.

| **Age** | **N**<br>(Total = 75) | **M age**<br>(months) | **SD age**<br>(months) | **Mean duration**<br>**of recording**<br>(in minutes) | **SD duration**<br>(in minutes) |
| --- | --- | --- | --- | --- | --- |
| **6-months** | 15<br>(6 females) | 6.43 | 0.48 | 25.09 | 6.81 |
| **9-months** | 15<br>(8 females) | 9.17 | 0.47 | 26.27 | 7.58 |
| **12-months** | 15<br>(7 females) | 12.47 | 0.99 | 25.78 | 4.62 |
| **24-months** | 16<br>(8 females) | 25.50 | 1.98 | 29.99 | 7.55 |
| **36-months** | 14<br>(8 females) | 36.16 | 2.86 | 32.51 | 7.22 |

#### Data acquisition and processing procedure

The experiment took place in a laboratory room with controlled lighting containing toys and objects; and a separate larger play area resembling a typical playroom with age-appropriate toys and books. There were three tasks: check-this-out game, spin-the-pots task, and free play. Participants wore a head-mounted eye tracker to record scene video and gaze points, and body sensors to record ECG and Acc. Further details are provided in the Supplemental Material.

#### Head-mounted eye tracker data recording and processing

Eye movements were recorded using a head-mounted eye tracker (Positive Science, New York, USA), which tracked the right eye at 30 Hz. Scene recordings were captured at 30 fps with 640×480 pixel resolution and a wide lens (W 81.88° × H 67.78° × D 95.30°). The Yarbus software (version 2.4.3, Positive Science) was used to map participants’ gaze points onto the scene video and calibrate the eye tracker offline, accounting for variations in eye morphology and the spatial location of fixated objects (Slone et al., 2018; Yu & Smith, 2013) (further details in the Supplemental Material). We calculated fixations from gaze points and labelled them using the GazeTag software (version 1, Positive Science). We only include fixations with duration > 100 ms. The labels represent 87 different items including toys, body parts, and other objects in the room. Approximately 10% of frames were unsuitable for labelling due to abrupt movements, participants removing the eye tracker, or technical errors.

#### Cardiac activity and body movement recording

ECG and Acc signals were recorded concurrently at 500 Hz using the Biosignalsplux device (PLUX Biosignals, Lisbon, Portugal). The ECG sensor’s three-electrode montage (including ground) was stuck to the left side of the chest, and the Acc sensor was placed at roughly the same location (Figure 2). The device also includes a light (Lux) sensor, enabling synchronisation of ECG/Acc data with the eye-tracking data. To ensure quick sensor placement, the Biosignalsplux hub and sensors were embedded in a custom-made shirt (Figure 2).

**Figure 2.**
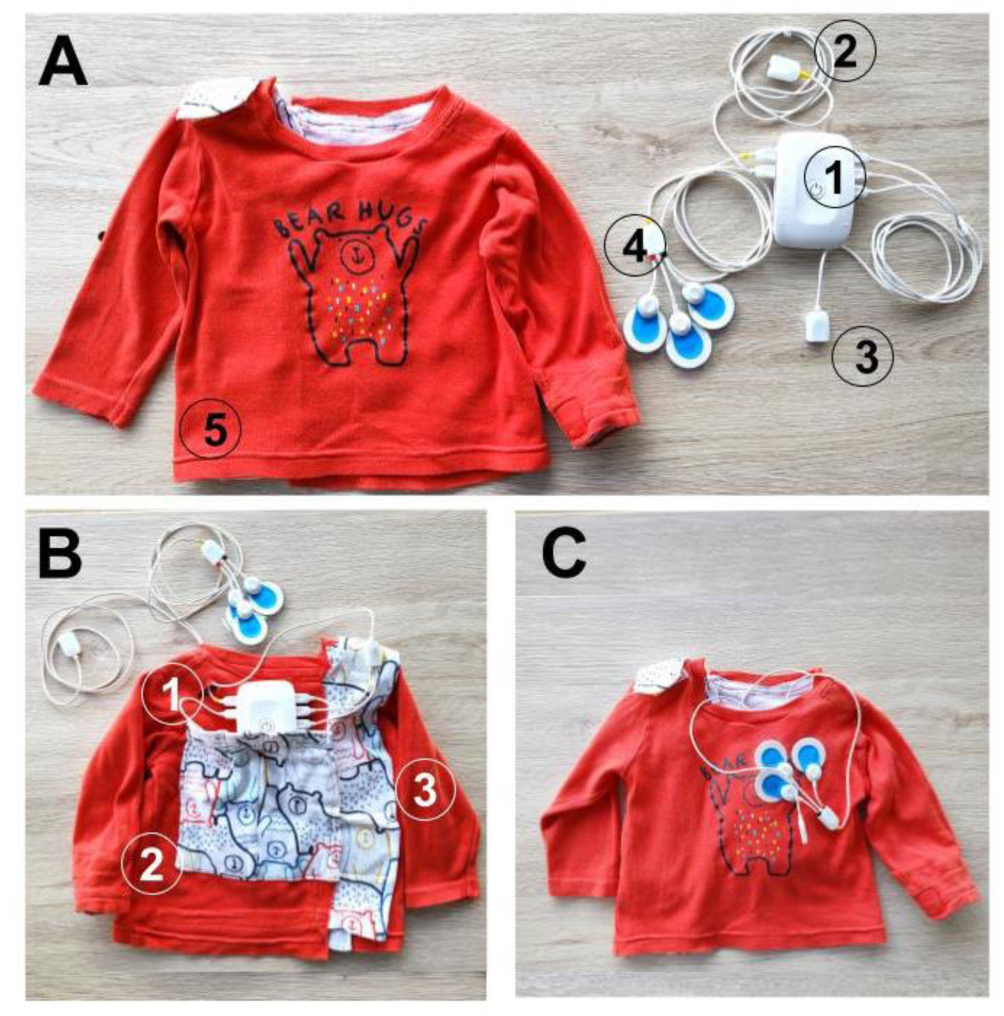
The wearable body sensor. (A) The Biosignals Plux wearable sensors: 1 - the data recording hub; 2 - the Lux sensor; 3 - the acceleration sensor; 4 - the ECG sensor with the Ambu blue electrodes (Ambu, Copenhagen, Denmark) attached; 5 - the custom-made shirt. (B) The back view of the shirt showing how the sensors were embedded: 1 - eyelet, which allows the ECG and Acc sensors to be brought from the back to the front and positioned on the left side of the chest; 2 - the back pocket, which holds the data recording hub; 3 - the shirt closes at the back via hook and loop. (C) Front view of the shirt with the sensors embedded, illustrating the location of the ECG sensor.

#### Sensor signal preprocessing and HR extraction

The triaxial accelerometer (Acc) data was centred by subtracting 2^15^ and normalised by dividing by 5000. The magnitude of acceleration was calculated as the L2-norm of the triaxial acceleration.

ECG processing and R-peak detection was handled using Python (Geangu et al., 2023; Mason et al., 2024), adapted from Neurokit2 code (Makowski et al., 2021). All R-peaks were then visually inspected; mislabelled peaks were manually corrected. The HR was calculated using the time interval between consecutive R-peaks (Δtpeaks): *HR (bpm) = 60/Δtpeak*s. The detailed procedure for noise correction has been described elsewhere (Mason et al., 2024). The parameters used in this study are in the Supplemental Material.

Bluetooth connectivity issues caused ECG/Acc data loss six times in total across all 75 participants, with a mean data loss duration of 159.1 s (*SD* = 106.5 s; range = 28.9 s - 279.5 s). No HR was calculated during these periods.

#### Human coding of sustained attention

We adapted the lab-based criteria for HRDSA for data acquired under naturalistic conditions. In particular, we defined SA as: (Criterion 1) occurring during periods with a deceleration in participants’ HR followed later in the period by an HR acceleration (Lansink et al., 2000; Lansink & Richards, 1997; Richards, 1987, 1997, 1998; Richards & Casey, 1991); and (Criterion 2) when participants fixate on and/or engage with a small number of objects during this time (Petrie Thomas et al., 2012). Detecting SA in naturalistic settings, such as free play, presents challenges compared to controlled lab conditions. Infants and toddlers in the lab have restricted movements and their HR is measured during a baseline period before an engaging stimulus is presented on a screen. However, free play lacks defined event structures and infants and toddlers can actively fixate and interact with objects, making it difficult to determine baseline HR measurements.

To address these challenges, we first developed a method to identify candidate periods corresponding to Criterion 1. We used a change point detection (CPD) algorithm to automatically detect abrupt decreases (SA onsets) and increases (SA terminations) in HR without the need for a pre-determined baseline (see *The Automatic Sustained Attention Prediction (ASAP) Model* section). This offers an objective approach to detecting abrupt changes by adaptively responding to the data, minimising the need for subjective, user-defined criteria, and holding the potential to accommodate variations attributable to individual differences, fluctuations in HR baseline, and the developmental change of HR. Second, we use eye-tracking data to determine whether these periods were associated with infants fixating on or following objects within the scene corresponding to Criterion 2.

#### Human coding protocol

Three experimenters (two authors) independently coded SA periods. Figure 3 illustrates the custom MATLAB graphical user interface for the protocol. First, detected change points in the HR time series were marked (Figure 3A, vertical dashed lines). Then the change in mean HR of a segment relative to the preceding segment was calculated.

**Figure 3.**
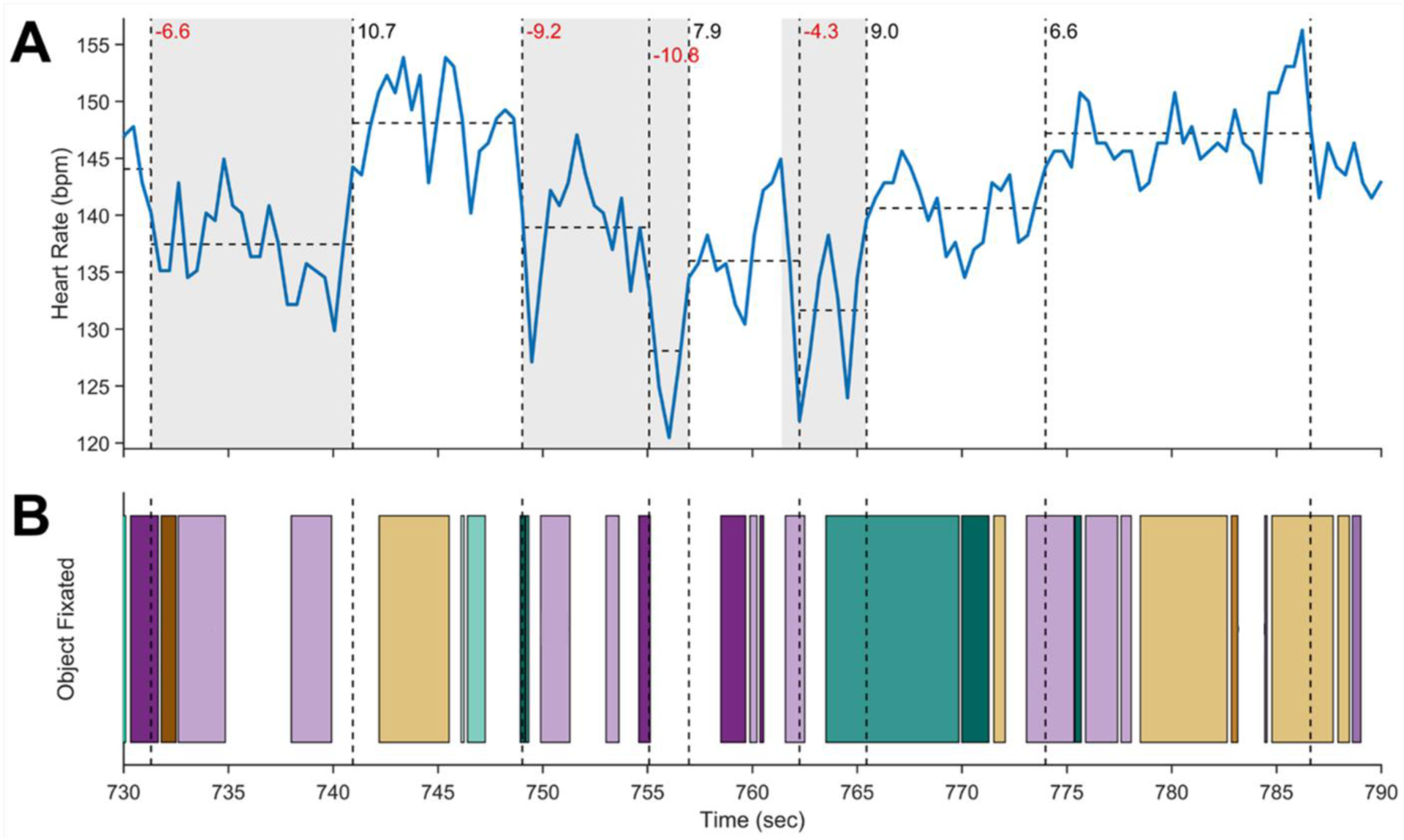
Example of human-coded sustained attention for a 6-month-old participant for a 1-minute time window. (A) The HR time series (in bpm) is the solid blue line. Change point timings are vertical dashed black lines which divide the signal into segments. The mean HR for each segment is shown with horizontal dashed black lines. The change in mean HR relative to the preceding segment is displayed in the top left of each segment (red indicates deceleration; black indicates acceleration). The grey regions indicate sustained attention periods. In the third grey region, the onset time was shifted to the first HR peak before the change point as the change in HR was −4.3 bpm, with the change between the HR peak immediately before and after the change point > 5 bpm. (B) The fixated objects from the eye-tracking data. Each unique colour bar represents a unique object (57 total objects), e.g., purple bars represent periods of fixation on a plush giraffe.

The change in mean HR on consecutive segments was next screened for putative SA periods as follows. The SA onset was set to the change point when there was a decrease of at least 5 bpm in the mean HR relative to the preceding segment. If the change was between 3 and 5 bpm, the coder assessed the peak-to-peak drop around the change point. If the drop exceeded 5 bpm, the onset was set to the time of the peak immediately before the change point (e.g., Figure 3A, third grey region). For consecutive drops meeting these criteria, the earliest drop determined the SA onset (e.g., Figure 3A, second grey region). The termination of SA was set at the change point with an HR increase of at least 5 bpm. For 3-5 bpm increases, termination was set to the nearest HR peak after the change point if the peak-to-peak rise was 5 bpm or more. SA periods shorter than 2 s were excluded (Petrie Thomas et al., 2012).

Lastly, coders assessed whether participants were looking at or following a small number of objects during the identified putative SA periods. This was done using the fixation-label time series (Figure 3B) and inspecting the screen video overlaid with the fixation crosshair. The experimental setup contained several toys or objects of potential interest (maximum 25), similar to a typical home (Anderson et al., 2022; Sun & Yoshida, 2022), and participants were free to move about the space. In most lab-based studies investigating HRDSA, complex and dynamic stimuli, which comprise multiple social and non-social elements and changes in scenes, were presented on computer screens (Bradshaw et al., 2024; Pérez-Edgar et al., 2010; Richards, 2010; Richards & Cronise, 2000; Richards & Gibson, 1997). In these studies, periods of HR deceleration indicative of SA have been associated with both brief and extended ‘looks’ towards the screen, each ‘look’ encompassing fixations towards different objects, as well as different scenes and events. Other lab-based studies presented infants with 1 to 6 toys, often handed to them one at a time (Luo & Franchak, 2020; Petrie Thomas et al., 2012; Yu & Smith, 2016). Considering the more naturalistic conditions in our study, we set the threshold to a maximum of 5 objects to be fixated or followed in order for a period of HR deceleration to be considered SA.

To assess inter-rater reliability, we randomly selected 15 participants from the five age groups, with two of the three coders independently coding each participant. Reliability was determined by counting overlapping attention and inattention periods between the two coders, allowing for any degree of overlap. The coders showed substantial agreement (Cohen’s κ: *M*=0.764; *SD*=0.116; range: 0.496 to 0.895).

#### Saliency and clutter extraction

We calculated saliency and clutter from the acquired scene video frames and fixation data during the naturalistic free-play periods. These measures can attract infant and adult participants’ attention (Amso et al., 2014; Hunter et al., 2023; Itti & Koch, 2001; Mahdi et al., 2018; Oakes et al., 2024; van Renswoude et al., 2019). The fixation distribution per age group is presented in the Supplemental Material (Figure S2, Table S1).

Figure 4 illustrates how we generated a saliency time series (see also Figure S3). Based on the duration of SA periods in our data, we first segmented the frames into consecutive 5-sec time windows to allow for the temporal integration of fixated visual information. Second, we created a 50-pixel radius circular mask (Bradshaw et al., 2023; Franchak et al., 2011; Kretch & Adolph, 2017) centred on each fixation and accumulated these masks within each window to create a binary fixation mask. On average, approximately six fixations contributed to each binary fixation mask, covering approximately 7% of the frame area (Table S2). Third, we extracted saliency maps from the frames, applied a binary fixation mask to each map within that window, and averaged the saliency values of pixels inside the mask to create the time series. For time windows with no fixations, the saliency/clutter values were removed from further analyses. We used the Graph-Based Visual Saliency (GBVS) algorithm to compute saliency (Harel et al., 2006) and the Feature Congestion measure to compute clutter (Rosenholtz et al., 2007). Full details of this procedure are provided in the Supplemental Material.

**Figure 4.**
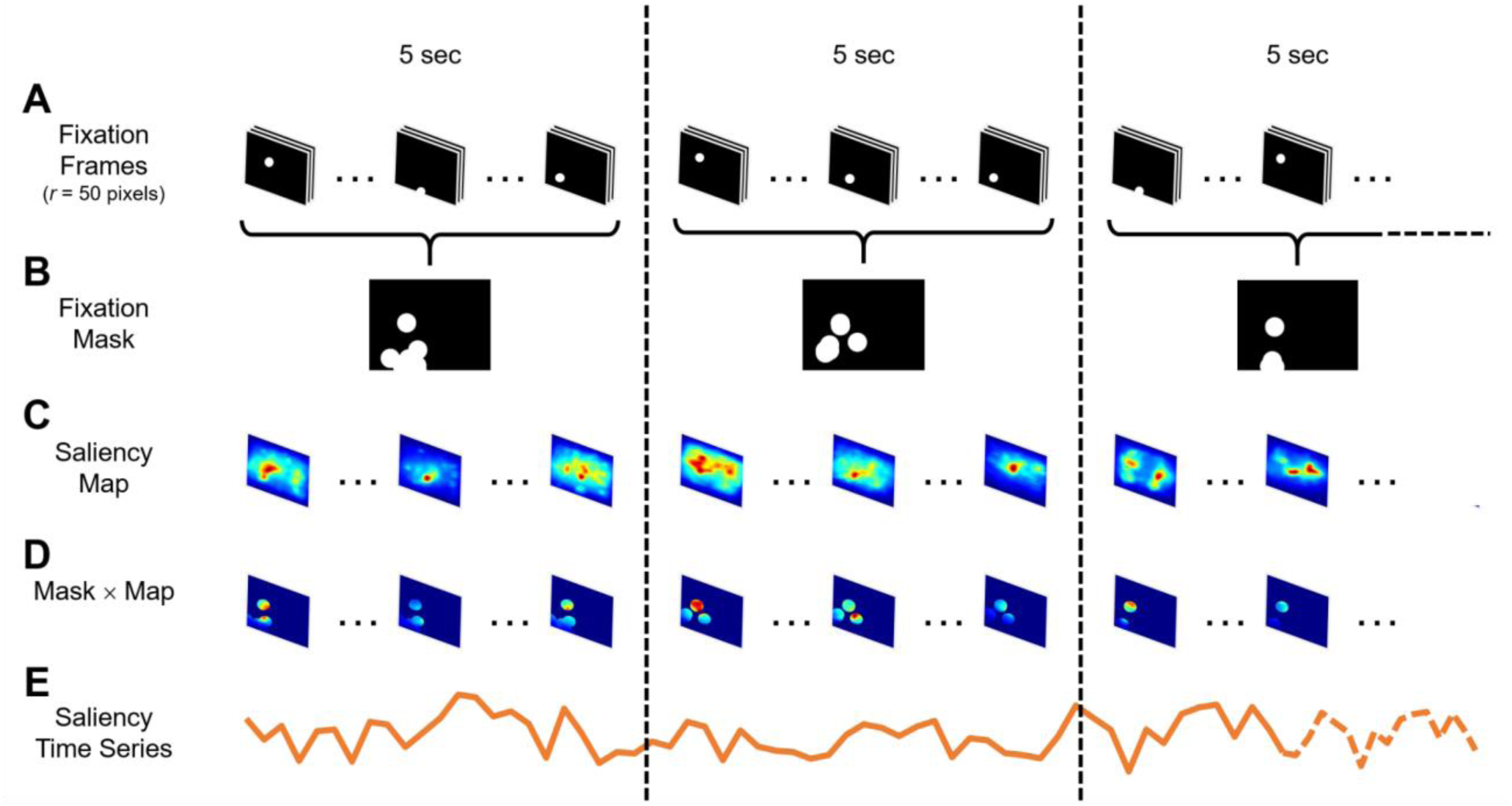
Example of a saliency time series during free play. (A) The fixation time series is divided into consecutive 5-sec time windows (150 frames at 30 Hz). A spatial window was created around each fixation (radius, *r* = 50 pixels; white circles). (B) All fixations within each 5-sec time window are accumulated to form a binary fixation mask (black pixels = 0, white pixels = 1). (C) The saliency maps within each window are extracted (15 frames at 3 Hz). (D) Each saliency map within a 5-sec time window is multiplied by the corresponding binary fixation mask for that window. (E) The saliency time series is created by averaging across all pixels within the respective binary fixation mask at each time point. The same procedure is used for the clutter time series. For a detailed illustration of the binary fixation mask applied to a saliency map, see Figure S3.

### The Automatic Sustained Attention Prediction (ASAP) Model

ASAP detected HRDSA by identifying abrupt changes in HR and leveraging information from HR and Acc signals associated with SA. Using the processed HR and Acc signals, we carried out feature extraction and feature selection to establish a classifier for attention prediction. As shown in Figure 5, the ASAP procedure consisted of three steps (colour-coded in Figure 5): *change point segmentation*, *point-wise classification*, and *segmentation refinement*.

**Figure 5.**
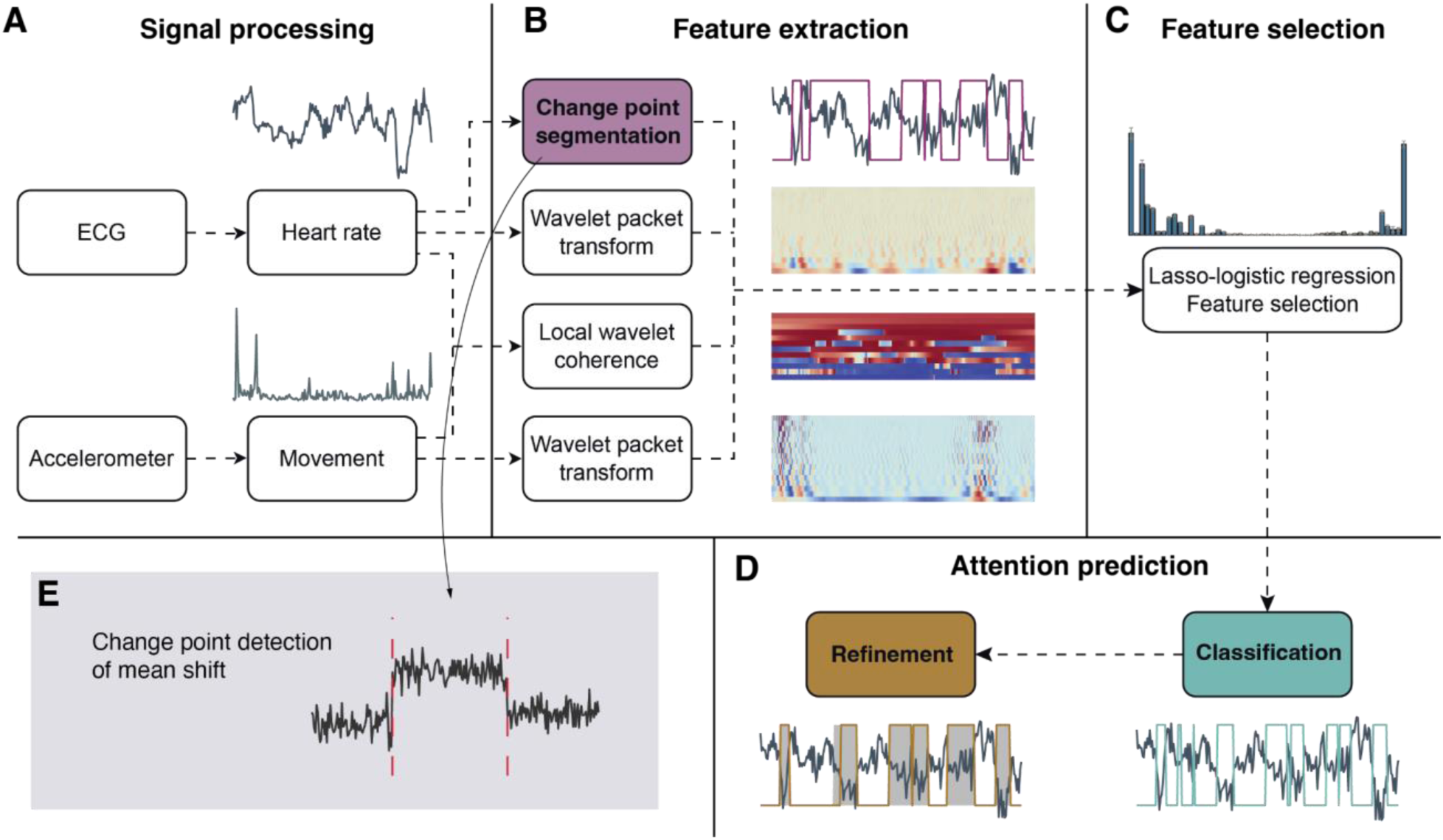
ASAP procedure overview. (A) Signal processing to extract heart rate (HR) and movement (acceleration magnitude). (B) Feature extraction to produce change point segments, wavelet packet transforms of HR and Acc, and local wavelet coherence between HR and Acc. (C) Feature selection using Lasso regularised logistic regression. (D) Attention prediction using logistic regression with selected features and further refinement to reconstruct the temporal structure. Coloured boxes in B-D indicate the three stages of attention prediction. (E) An example of change point detection of mean shift.

#### Step 1: Change point segmentation

Change point detection (CPD) is a statistical method that detects changes in properties (e.g., mean, variance, slope) in a time series. Here, the change point segmentation step (Figure 3) identified boundaries of putative SA periods indicated by steep decreases and increases in the mean of HR (Figure 5E). Given that infant free play is characterised by frequent short periods of SA and that the heartbeat defines a fundamental minimum resolution of 2-3 samples per second for observing changes, we adopted a CPD methodology that produces change points best fitting these resolution constraints: wild binary segmentation (WBS2) with a model selection criterion of the steepest drop to low levels (SDLL) (Fryzlewicz, 2020) (for the selection motivation, see Supplemental Material). The method is available from CRAN via the R package *‘breakfast’* (version 2.2), using *‘wbs2’* and *‘sdll’* options.

Applied to our dataset, this approach had an average rate of CPD of one change point per 8 to 9 heartbeats. Change points were classified as descending and ascending by calculating the local slopes of HR change. We then applied the same rules as for the *human coding protocol* to screen and refine the boundaries of SA. Step 1 resulted in a binary time series indicating putative attention and inattention segments.

#### Step 2: Point-wise classification

*Feature extraction.* Time points were classified mainly based on the temporal and spectral information extracted from HR and Acc and the putative SA segments identified in Step 1. 51 predictor features were extracted (see Table S3 and Figure 6 for full list). (1) The HR and Acc magnitude. (2) Time-frequency decomposition using the wavelet packet transform was used to analyse HR and Acc over different time scales (MATLAB 2022a routine ‘*modwpt’*, Daubechies wavelet with two vanishing moments), yielding 16 features (frequency bands) for each measure (WPT-HR and WPT-Acc). (3) The time-evolving spectral cross-dependence between HR and Acc was estimated using multivariate locally stationary wavelet processes, with the localised coherence Fisher’s-z transformed (R package *‘mvLSW’* version 1.2.5) (Park et al., 2018), yielding 12 features (LSW-HR-Acc). (4) HRV measures from the recording session were included, specifically the standard deviation of inter-beat intervals (SDRR) – the time deviation between consecutive R-peaks. (5) A binary time series derived from the CPD step was included, indicating potential attention and inattention states (Change point binary). (6) Segment duration (Duration) and time to the previous segment (Latency) yielded two more features. (7) Lastly, age in months was included.

**Figure 6.**
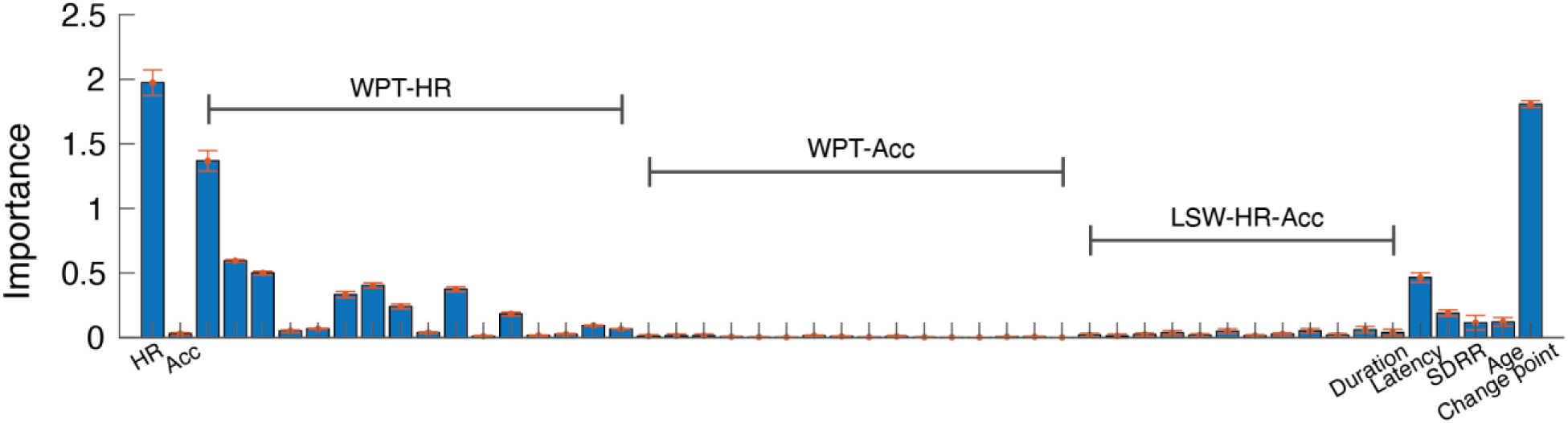
Feature importance by Lasso regularised logistic regression. Importance is measured by the absolute values of the standardised logistic regression coefficients, averaged across the 5-fold cross-validation. Error bars are standard deviations across the 5-fold cross-validation. Selected features had mean Importance > 0.03 (threshold determined by cross-validation). WPT-HR: wavelet packet transform of HR; WPT-Acc: wavelet packet transform of Acc; LSW-HR-Acc: local stationary wavelet estimated coherence between HR and Acc. In these cases, each bar corresponds to a frequency band.

These variables formed the predictors, *X*, in the model *Y = f(X)*, where *Y* represents a binary time series of attention and inattention from human coders. We initially included the above features as predictors, and then performed feature selection to determine the most informative ones to predict attention. All variables were sampled at 2 Hz.

#### Feature selection

The 75 participant sessions were split into training and test sets (∼9:1), where each partition contained randomly selected sessions with approximately equal age distribution. The training set consisted of 67 participant sessions (194360 time points), and the test set consisted of 8 participant sessions (22367 time points). Our model selection procedure was carried out in the training set. Feature selection was performed using regularised logistic regression, which shrinks a subset of the estimated coefficients to zero by employing an L1-norm penalty (Lasso) for covariates deemed to have non-significant contributions to attention (Tibshirani, 1996) (see Supplemental Material for the mathematical formulation). The tuning parameter (λ) that controls the strength of the L1-norm penalization was determined by cross-validation (λ = 3.4*e* − 5). The regularised logistic regression model was trained using the ‘*lassoglm*’ MATLAB routine with 100% L1 penalty.

The Lasso approach preserved 24 out of the 51 feature variables (Figure 6 and Table S3), including the HR, a subset of wavelet transforms of the HR (WPT-HR in Figure 6; 13/16 selected; see Table S3 for selected frequency bands), duration, latency, SDRR, change point binary, and age. The Acc alone and its wavelet transform (WPT-Acc) were determined to be unimportant, but five of the twelve HR-Acc coherence bands (LSW-HR-Acc) were included. The logistic regression model was then retrained without regularisation using the training set based on the 24 features. The performance was then tested on the test set. Using other classifiers resulted in comparable performance (Figure S4, Table S4).

#### Model training

Imbalanced class distribution, where one class significantly outnumbers the other, can bias model training. In our case, attention occupied about 29% of the total time. To address this, we tested whether balancing class distribution could improve model performance using 5-fold cross-validation. Data balancing was achieved using the Synthetic Minority Over-sampling Technique (SMOTE) (Chawla et al., 2002). SMOTE improved the model performance based on the F1-score criterion. We showed that the model’s performance remained stable regardless of dataset size, with performance metrics plateauing when trained on more than half of the training set (see Supplemental Material for sensitivity test; Figure S5). To optimise model accuracy for deployment with future data collected in natural environments, we provided the final model trained on the entire dataset with SMOTE oversampling applied.

#### Step 3: Segmentation refinement

Point-wise classification (Step 2) did not preserve the temporal structure of the putative SA segments obtained in Step 1, and could disrupt long SA periods. To maintain the temporal structure, we merged adjacent segments based on the information from Step 1 and Step 2. First, segments identified in Step 2 that were 2.5 seconds or less apart were merged. Next, if the predicted attention in Step 2 covered more than 90% of a Step 1 SA segment, the entire segment was classified as an SA period, provided that no attention span exceeded 50 seconds. These criteria were established using 5-fold cross-validation within the training set, which optimised on producing an attention duration distribution similar to the human-annotated distribution. The Kolmogorov-Smirnov (KS) test was applied to compare the model distribution to the human distribution.

#### Model evaluation

The performance of the ASAP model was evaluated on the test set (8 sessions from all age groups of 6-, 9-, 12-, 24-, and 36-months). First, we assessed the point-wise performance of the method in comparison to human coding for different metrics: 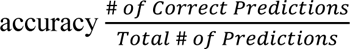, 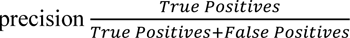, 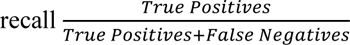, 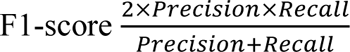, and inter-rater reliability (Cohen’s κ) at each step. Second, the KS test was used to assess the similarity of the predicted duration distribution compared to the human-coded data. Finally, we examined whether our prediction method could preserve the natural statistics associated with attention as identified by human coding (*Model application*). The criterion of statistical significance was set to 0.05.

## Results

### Model performance

The ASAP procedure progressively approximated human-coded attention periods through the three steps (Figure 7A, B). The CP segmentation phase (Step 1) attained a point-wise accuracy of 80±5% (mean±SD), precision of 60±10%, recall of 92±6%, F1-score of 0.72±0.07 and Cohen’s κ of 0.76±0.07. An all-negative prediction, based on the assumption that participants were not paying attention for the majority of the time, resulted in a significantly lower accuracy of 71%. Conversely, an all-positive prediction led to a precision of 29%, consequently producing a null F1-score of 0.45. The failure to achieve 100% recall can be attributed to adjustments made during the human coding process incorporating the video and fixation data. Additionally, the duration of attention segments exhibited a broader distribution than the human-coded distribution (*p* = 0.0056, KS test, Figure 7C).

**Figure 7.**
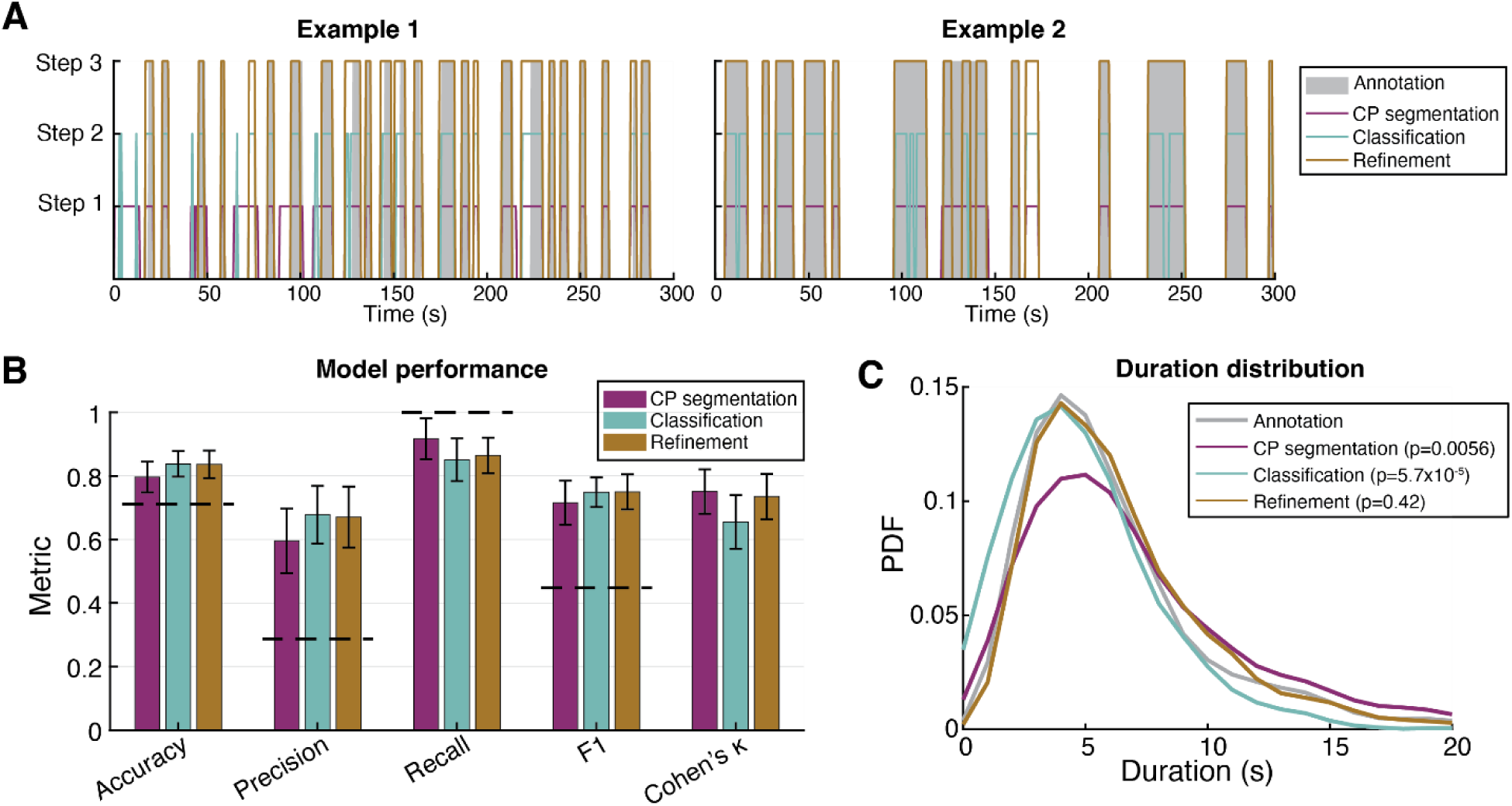
ASAP performance. (A) Two examples of attention prediction through 3 steps. (B) A summary of performance metrics. Colour codes are identical to (A) and Figure 5. Dashed lines indicate null estimates. The null is obtained for accuracy when guessing all negative; for precision, recall, and F1, the null was calculated with all-positive guesses. Error bars are standard deviations. (C) Distributions of segment duration were obtained from each step. P-values revealing the distribution similarity are based on KS test. CP: change point. PDF: probability density function.

The classification step (Step 2) improved the prediction accuracy to 84±4% (*p* = 0.0060, paired *t*-test between Step 1 and 2) by increasing precision to 68±9% (*p* = 1.9e-4, paired *t*-test). This step reduced recall to 85±7% (*p* = 0.0025, paired *t*-test), indicating a trade-off between precision and recall. Cohen’s κ also dropped to 0.66±0.08 (*p* = 5.0e-4, paired *t-test*) due to the reduction in recall. Step 2 fragmented long attention periods, resulting in a duration distribution skewed towards shorter durations (*p* = 5.7e-5, KS test, Figure 7C). Overall, Step 2 improved the accuracy of point-wise predictions but did not maintain the temporal structure.

The refinement phase (Step 3) merged short segments to reconstruct long attention spans. The point-wise performance remained largely unchanged, with accuracy of 84±4%, precision of 67±10%, recall of 86±6%, F1-score of 0.75±0.06, and Cohen’s κ of 0.74±0.07. Importantly, this phase restored the duration distribution to be statistically comparable to the human-coded distribution (*p* = 0.42, KS test, Figure 7C). Therefore, Step 3 was crucial for recovering intact attention periods. Additionally, we did not observe a significant effect of age on any of the metrics at any step (*p*s > 0.2, linear regression).

### Model application: Cross-validation of the ASAP model with egocentric visual information

An infant’s attentional state is linked to their sensory experiences within natural environments. We estimated the changes to visual saliency and clutter of fixated regions over time (see *Saliency and clutter extraction* and Figure 4) and compared these measurements between ASAP and human-coded attention and inattention periods, while also examining potential developmental changes. Only data from free play sessions with at least one minute of cumulative recorded fixation (N=72; 966±368 s) were included in this analysis. We standardised saliency and clutter measures within each session to eliminate systematic variation across participants. The session means of each measure were analysed using a linear model (*fitglm* in MATLAB, 2022a), with three factors: attentional state (attention vs. inattention), data source (human coding vs. ASAP prediction), and age, along with their four two-way interactions and one three-way interaction term (see Supplemental Material, Table S5). A Bonferroni correction was applied to adjust for multiple comparisons across factors, interactions, and responses.

We found the attentional state had a significant effect on mean saliency, with higher saliency of fixated regions during attention than inattention periods (Bonferroni-corrected *p* = 9.2e-7; Figure 8A). There was also a significant interaction between attentional state and age for saliency (Bonferroni-corrected *p* = 0.0011; Figure 8B): the saliency of fixated regions decreased with age during attention periods but remained relatively constant during inattention periods. No significant effects or interactions involving the data source were found (complete statistical results in Table S5). Additionally, no attentional or age-related effects on the visual clutter of fixated regions were observed (all *p*s > 0.05; Figure 8C, D; Table S5).

**Figure 8.**
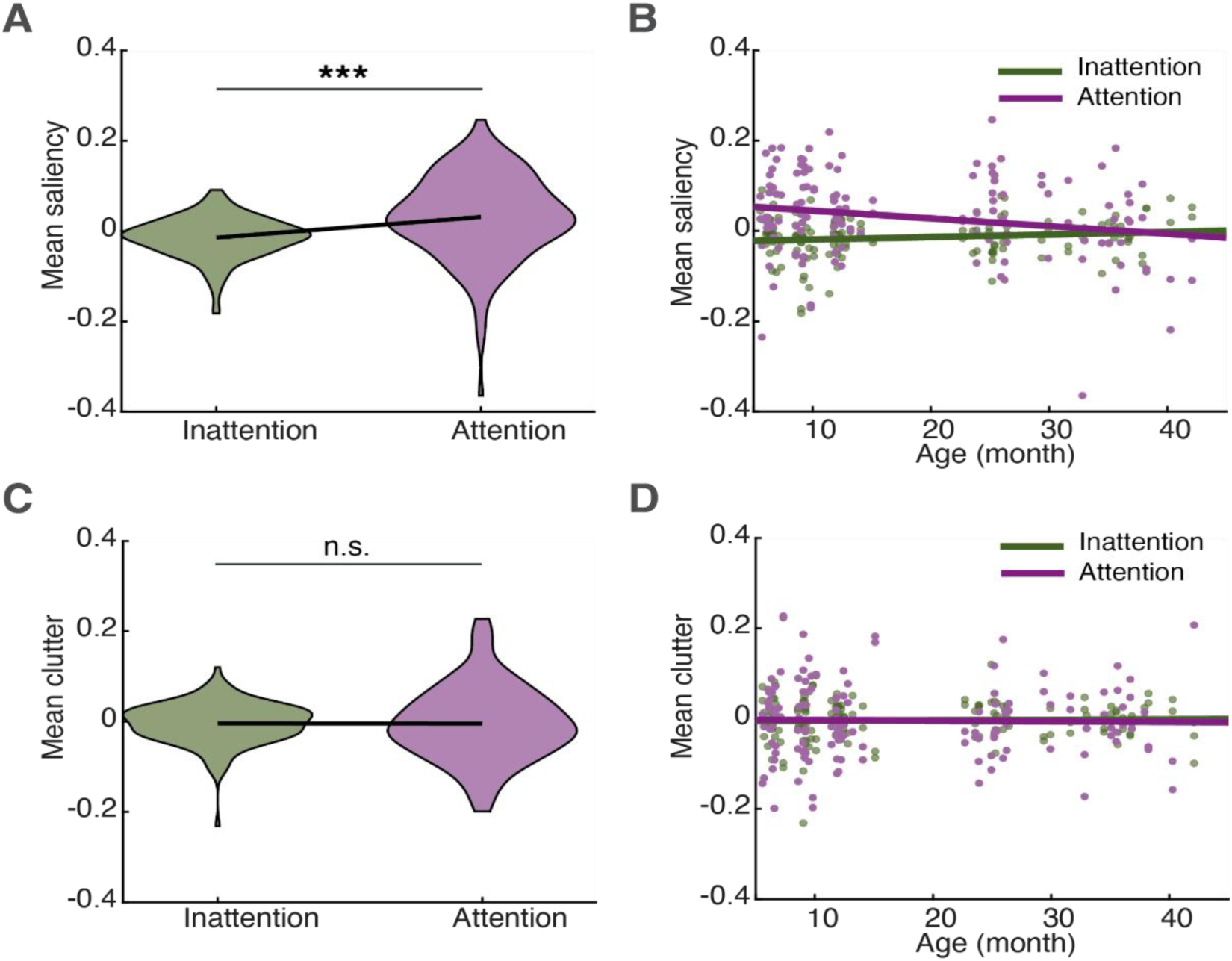
Effects of attention on visual features. (A) Violin plot of the attentional effect on visual saliency. (B) Interaction effect on visual saliency between attentional states and age. Lines are fitted by linear regression. (C) The same as (A) for clutter. (D) The same as (B) for clutter.

## Discussion

There has been increasing interest in employing naturalistic approaches to study human development (Dahl, 2016; Smith et al., 2018; Smith & Karmazyn-Raz, 2022; Sparks et al., 2024; Sullivan et al., 2021). Sustained attention (SA), a cognitive ability that emerges during infancy, is particularly important due to its widespread implications across various developmental domains (Brandes-Aitken et al., 2023; Reynolds & Romano, 2016; Ruff & Lawson, 1990). Despite its significance, most current SA developmental research relies on lab-based paradigms, with limited understanding of how mutual interactions between the developing infant and their everyday environment contribute to its emergence. In this study, we propose an innovative algorithm - ASAP - that harnesses signals recorded unobtrusively from wearable technology (e.g., the EgoActive platform (Geangu et al., 2023)) to detect infant SA manifested spontaneously in the natural environment.

The existent lab-based research demonstrates that heart rate (HR) can provide rich information about the ANS’ regulation of homeostasis, physiological arousal and cognitive states, including SA (Forte et al., 2019; Richards & Casey, 1991; Thayer et al., 2009). ASAP was developed by acquiring a dataset of multimodal signals (ECG, Acc, and eye-tracking) in 6- to 36-month-olds during different naturalistic situations, using this multimodal signal to code periods of SA informed by the empirical evidence (Richards & Casey, 1991), and finally combining this “ground truth” with informative features extracted from the temporal and spectral properties of/between the HR and Acc signals to train the classification model.

The key HR features contributing to successful SA detection are the HR deceleration during periods of SA relative to those of non-attention (Richards & Casey, 1991), and the HR fluctuations in both time and frequency domains (Addison, 2005; Akselrod et al., 1981; Börger et al., 1999; Borjon et al., 2016; Hansen et al., 2003; Pichot et al., 1999; Toledo et al., 2003; Van Roon et al., 2004; Zhang et al., 2022). The feature importance analysis revealed that HR fluctuations in the 0.03-0.15 Hz and 0.3-0.6 Hz frequency bands contributed significantly to predicting SA (Figure 6 and Table S3), corresponding to the two distinct peaks previously observed in the infant HR power spectrum (Finley & Nugent, 1995). A reduction in body movement has also been previously used as a criterion for determining attention (Byrne & Smith-Martel, 1987; Chen et al., 2016), and associations have been shown between changes in HR and body movement linked to measures of attention (Bazhenova et al., 2003; Porges et al., 2007; Wass et al., 2016; Wass et al., 2015). However, our feature selection did not identify the Acc magnitude as playing a significant role. This may partly be due to using Acc magnitude of the torso, rather than the movement of the head or limbs as in other studies (Wass et al., 2016; Wass et al., 2015). Our choice for relying on the torso movement was largely determined by the fact that in the wearable sensors typically used for naturalistic recordings, accelerometers are often placed on the torso (Geangu et al., 2023). Furthermore, in nonhuman primates, torso movement was found to be more coupled with HR than limb movement (Zhang et al., 2022). Although Acc magnitude did not predict SA in this study, its coupling with cardiac activity did, especially at very slow frequencies below 0.1 Hz (Table S3). This frequency range aligns with previously reported spontaneous brain fluctuations that are associated with arousal and its coordination with movement in humans and nonhuman primates (Gutierrez-Barragan et al., 2019; Raut et al., 2021; Zhang et al., 2022). Considering that some of the key features in defining periods of SA are represented by the deceleration and acceleration of the HR, as well as HR fluctuations in the frequency domain, our findings suggest that these are less likely to be the direct result of the changes in body movement.

We also investigated whether the variability in performance across sessions was driven by intrinsic dataset characteristics or biases inherent to the model. By analysing the correlation between human coder agreement and the agreement between human coders and the ASAP model, we found that intrinsic factors influenced performance for both (Supplemental Material, Figure S6A, B). Further analysis indicated that SDRR (an HRV measure) was a key intrinsic feature that contributed to SA classification performance (Figure S6C). Previous studies have shown that respiratory sinus arrhythmia (RSA), a separate HRV measure within the respiratory frequency range, predicts the extent of HR deceleration entering the attentional phase (Richards, 1987). Individuals with higher RSA levels (also higher SDRR) are more resistant to distraction (Hansen et al., 2003; Richards, 1987). Therefore, higher HR variation may lead to more distinct HRDSA, potentially reducing the confusion in the attention classification task. This correlation supported the inclusion of SDRR as a feature in the attention prediction model, and our feature selection analysis further validated its relevance.

We applied ASAP-labelled data to study visual attention development by leveraging the egocentric video and fixation data and generating the perceptual features of visual saliency and clutter. Three key findings emerged. First, and of critical importance, the results were similar for the periods of attention determined by human coders and those detected by ASAP (Table S5). This demonstrates that implementing ASAP can be a less costly (time and human resources) approach, making it ideal for studies involving large data recorded in the natural environment or in the lab. Second, similar to some previous studies (Pomaranski et al., 2021), during SA periods, infants fixated on areas with higher perceptual salience than during periods of inattention. Older infants were observed to fixate on less perceptually salient regions during sustained attention than younger infants. This could reflect the older infants’ increased reliance on more high-level properties of the scenes, although further consideration of other factors (e.g. local meaning) that tend to covary with salience would be required to more definitively conclude this effect (Oakes et al., 2024). Third, unlike some of the previous studies (Helo et al., 2017), the areas fixated during attention and inattention did not seem to differ in terms of visual clutter, irrespective of infants’ age. In part, this could be due to the fact that here we considered the amount of fixated feature congestion, which may be less relevant for allocating and sustaining attention compared to how cluttered the entire scene is (Helo et al., 2017). Our results help to replicate and extend findings from lab-based studies under constrained conditions to more naturalistic scenarios. Importantly, for the first time, the relation with visual saliency is reported for HRDSA, supporting the interpretation that visual saliency not only influences what infants are likely to orient towards but also what is likely to be processed in depth (Lansink & Richards, 1997; Ruff & Rothbart, 2001).

### Limitations and future work

ASAP is developed based on HR-defined visual attention. Our model yields a weaker precision (∼67%) relative to the other metrics, which may be partly due to other events inducing similar HR changes. Attention in other sensory modalities (e.g., audition) can also be accompanied by HR deceleration (Graham & Jackson, 1970), resulting in an HR waveform similar to that observed during visual attention. Additionally, vocalisations have been associated with an HR dynamic that mirrors the one observed during visual SA (Borjon et al., 2016; Zhang et al., 2022). Other information can help differentiate these possibilities from visual SA (the ground truth in our study), and future work could extend ASAP to detect sustained auditory attention while also incorporating infants’ and toddlers’ vocalisation as features, potentially improving its performance.

## Conclusion

We demonstrated that ASAP achieved solid performance in the challenging task of detecting periods of sustained attention based on HR and Acc signals across the age range and conditions considered here, including natural, free-play behaviours. This is critical as HR and Acc are the predominant signals recorded with unobtrusive wearable sensors for infants and are invariant across platforms. Although future work can expand ASAP to include other sensory modalities and increase its performance metrics, it can currently provide a powerful tool to detect visual sustained attention in natural settings (e.g., home). This reduces the reliance on fixation-based methods, which require eye trackers that are not suitable for long unsupervised recordings with infants and young children. Together with available wearable technology, ASAP provides unprecedented opportunities for understanding how attention develops in the context of infants’ and toddlers’ daily experiences, and to support explanations of attention development with high ecological validity.

## Author Contributions

Conceptualization: EG, PMC, YZ, HM, MIK, QCV Methodology: EG, PMC, YZ, QCV, HM, MIK, MCG, DM

Formal Analysis: YZ, QCV, HM, MIK, PMC, EG Investigation: EG, PMC, MCG

Writing – Original Draft Preparation: EG, PMC, YZ, HM, MIK, QCV Software: YZ, HM, QCV, DM

Validation: EG, PMC, YZ, HM, QV

Resources: EG, QCV, MIK

Data Curation: EG, QCV, HM, PMC

Visualisation: YZ, QCV, HM, EG

Supervision: EG, QCV, MIK

Project Administration: EG

Funding Acquisition: EG, QCV, MIK

## Conflicts of Interest

The author(s) declare that there were no conflicts of interest with respect to the authorship or the publication of this article.

## Acknowledgment

We would like to express our gratitude to all families who dedicated their time to participate in this research. Without their continued interest in our research and desire to help, these findings would have not been possible. The authors would also like to thank Brigita Ceponyte, Emily Clayton, Nicoleta Gavrila, Aastha Mishra, Lauren Charters, Fiona Frame, and Laura Tissiman for their invaluable support and help throughout the project.

## Funding

The work presented in this paper was partially funded by Wellcome Leap - The 1kD Program.

## Supplemental Material

### 1. Tasks description

The experimental session was divided into two parts. Part 1 consisted of the initial setup, synchronisation of the devices, and the eye-tracking calibration procedures, and took place in the laboratory room. The experimenters first fitted the eye tracker and body sensor on the participant. The recordings from the eye tracker and body sensor were then temporally synchronised by turning on and off the room light. Once the participants were at ease, *peek-a-boo* and follow-the-toy games were used to collect calibration points for the offline calibration of the eye-tracker. Five calibration points were collected as part of the *peek-a-boo* game. Using a blackboard with five cutout windows (located in the four corners and centre), the experimenter showed and hid a red dinosaur hand puppet and tried to draw participant’s gaze towards the puppet. The puppet was randomly shown several times at the five spatial locations (Kretch et al., 2014). The participants sat approximately 3.5 m from the blackboard throughout this step. During the *follow-the-toy* game, the experimenter sat close to the participant and directed their gaze to a series of colourful squeaky toys located at hand level (Yu & Smith, 2017) and other points covering space along the horizontal (azimuth), vertical (elevation) and depth axes not otherwise captured during the *peek-a-boo* game.

Part 2 consisted of three tasks aimed at eliciting visual attention in a variety of situations. It included more structured situations (the *check-this-out* game and the *spin-the-pots* task) and more naturalistic scenarios (free play). All participants were involved in the *check-this-out* game and free play. Since the *spin-the-pots* task was designed for older infants, only 24- and 36-month-old infants participated. The *check-this-out* game consisted of a period of semi-structured play. Parents and participants were invited into a play area and asked to find a comfortable position near the centre of a play mat, similar to what they would typically do at home for play. One of the experimenters then pointed to and showed a series of six target toys strategically arranged amongst other toys on a shelf located about 3.5 m in front of the child. The experimenter showed the target toys twice in the same sequential order to all participants while trying to capture their interest. The toys and their locations were consistent across all participants. This task lasted 91 s on average (*SD =* 25 s).

The *spin-the-pots* task tests working memory (Hughes & Ensor, 2007; Zimmermann et al., 2021). In this task, children must find six stickers hidden under eight pots of different colours. The location of the pots is hidden from view and rotated 180 degrees after each trial. During each trial, the child can only open one pot from the eight available locations. An error occurs when the child reaches for an empty pot. This task lasted 415 s on average (*SD =* 130 s).

For free play, parents and participants could play with the toys available in the play area, move around, and/or interact with each other as they would typically do at home. This unstructured playtime lasted an average of 1120 s (SD = 359 s). During unstructured play the collection of toys was the same for all participants. However, each child-parent dyad was found to differ in their selection based on their interests.

### 2. Head-mounted eye-tracker calibration protocol

We followed an offline calibration protocol recommended for naturalistic tasks (Slone et al., 2018; Yu & Smith, 2013), using the 15 calibration points acquired at the beginning of the experimental procedure (see Supplemental Material – section 1). After the experimental session, these calibration points were used to fit the corresponding coordinates in every frame taken by the scene camera. We implemented a quality threshold of >0.80 to ensure a good fit estimation in the eye-to-scene calibration, as recommended by Yarbus software.

### 3. ECG signal preprocessing and HR extraction

Initial ECG processing followed the pipeline laid out in Mason et al., (2024. This is based on using the Neurokit2 Python package (Makowski et al., 2021) default method with an additional 15 Hz high-pass frequency filter in preprocessing and local peak correction after. No square wave noise was exhibited by the Biosignalsplux sensor, so no square wave filter was used. After processing and peak detection, the initial list of peaks was parsed. Peaks were then shifted/added/subtracted by hand in order to form an accurate final list of peaks.

HR processing was done according to the protocol for clean HR datasets described in (Mason et al., 2024). Specifically, a local median filter of width 11 with an activation threshold of 1.7x the median was used to detect outliers in the data (including the gaps in time series caused by Bluetooth dropout). The HR value within any detected gap (median duration: 1.2s) was filled in by linearly interpolating between the intervals of the nearest time point on either side of the gap that fell within the threshold. Signal quality index analysis (Mason et al., 2024) was found to have minimal effect on the processing due to the high quality of the peak detection, and so was not used.

### 4. Fixation Distribution During Play Period

**Figure S2.**
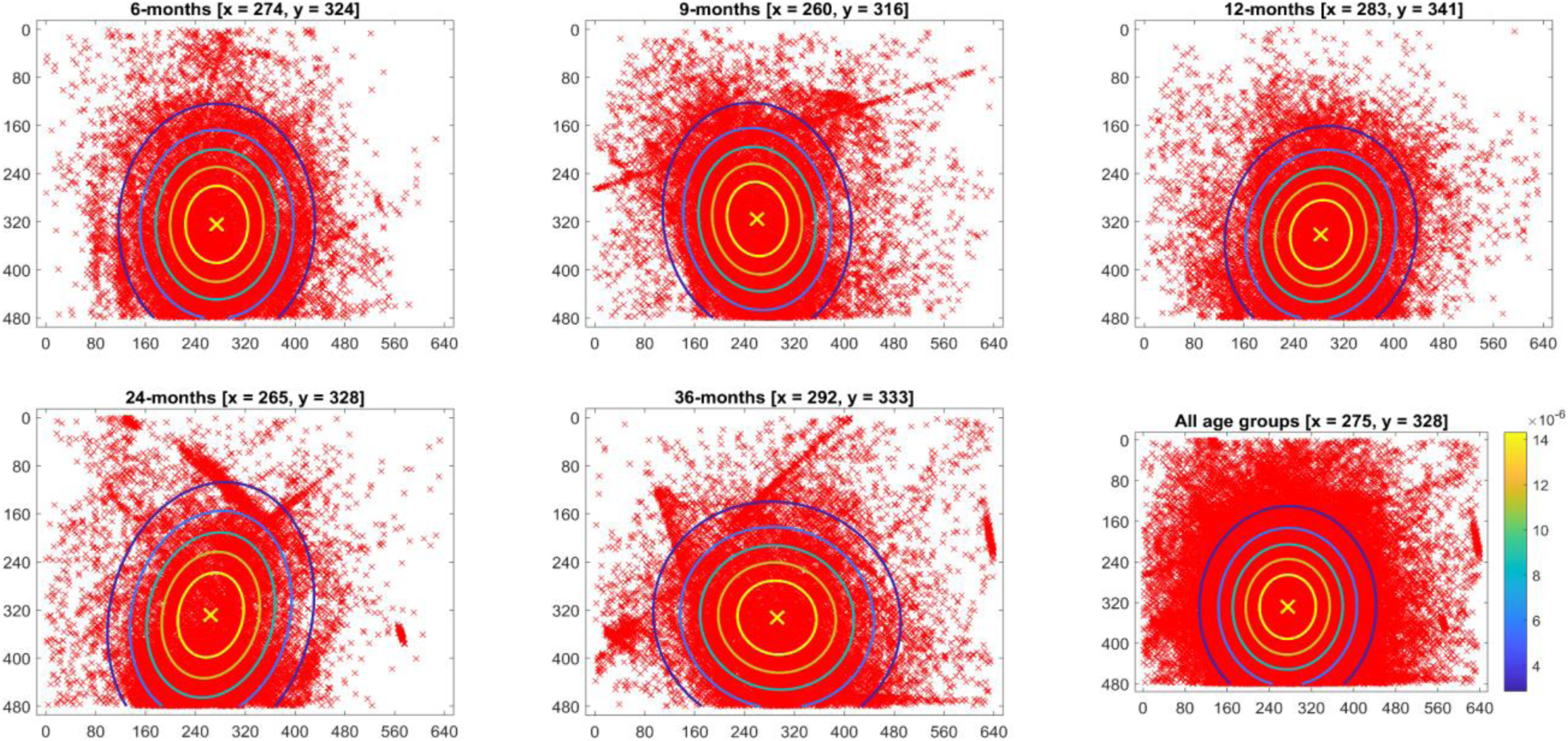
Fixation distribution as a function of age group. Red x’s represent individual fixations, and the yellow x represents the mean fixation (indicated by the *x*- and *y*-coordinate in plot title). A multivariate normal probability density function was fitted to each distribution, with the means and covariance matrix of the *x*- and *y*-coordinates. The contours represent equal-height isolines (5 levels) of the fitted probability density function (colour bar), with fixations more likely towards the mean fixation coordinates (yellow x and contour). See Table S1 below for additional statistics.

**Table S1.** Fixation distribution as a function of age. The *x*- and *y*-coordinates are in pixels (frame size: 640 x 480 pixels). Play duration is in seconds. The standard deviation (SD) is in brackets. The cov(*X*, *Y*) is the covariance between the *x*- and *y*-coordinates across all fixations.

| Age Group | Total #<br>Fixations | Fixation Distribution |  |  | Mean Play<br>Duration<br>(SD) |
| --- | --- | --- | --- | --- | --- |
|  |  | Mean X<br>(SD) | Mean Y<br>(SD) | cov(X,Y) |  |
| 6-months<br>(N = 15) | 17881 | 274.3 (83.3) | 324.4<br>(106.1) | -59.9 | 971.1 (335.7) |
| 9-months<br>(N = 15) | 23087 | 260.1 (82.7) | 315.5<br>(102.1) | 581.0 | 1223.9 (365.5) |
| 12-months<br>(N = 15) | 21486 | 283.0 (81.4) | 341.4 (95.1) | -614.7 | 1168.9 (254.6) |
| 24-months<br>(N = 16) | 18600 | 264.8 (87.5) | 328.4<br>(116.7) | -1439.6 | 1042.1 (415.2) |
| 36-months<br>(N = 14) | 21223 | 292.1<br>(104.8) | 332.7<br>(101.8) | 470.1 | 1203.9 (377.4) |
| All age groups<br>(N = 75) | 102277 | 274.9 (88.7) | 328.4<br>(104.5) | -92.0 | 1119.8 (358.7) |

### 5. Saliency and clutter extraction

We created 1-dimensional saliency and clutter time series based on fixated regions for each participant and investigated the extent to which attention periods predicted these time series. Here, we detail the procedure for visual saliency. The same procedure was used to generate the clutter time series. The fixation data consisted of a time series (frames) of *x*- and *y*-coordinates as computed by the Positive Science software. First, for each fixation, we created a circular mask centred on the fixation with a radius, *r* = 50 pixels. This radius reflects the precision of the eye tracker and allows for spatial summation of visual information around the fixation. The radius is based on previous studies that used screen-based studies with the same system (Bradshaw et al., 2023; Franchak et al., 2011; Kretch & Adolph, 2017). We then segmented the fixation frames into consecutive 5-sec windows. We used a temporal window to reflect how infants and toddlers may integrate visual information at fixation across a short temporal period. The value was partly based on the duration of attention periods we observed in the data. Second, all circular masks within each time window were combined to create a binary fixation mask (see Table S2 for the characteristics of binary fixation masks as a function of age). Third, we extracted saliency maps from the video frames. For each 5-second time window, we multiplied each saliency map in the window by the same fixation mask from that window (see Figure S3 for an example of a binary fixation mask applied to a video frame and saliency map). Finally, we averaged the saliency across all pixels within the fixation mask to create a 1-dimensional saliency time series. There were 5-second time windows with no fixations; in these cases, the saliency/clutter values were removed from further analyses.

We used the Graph-Based Visual Saliency (GBVS) algorithm to compute saliency maps (Harel et al., 2006), as evidence suggests that this algorithm predicts infant and adult fixations better than other saliency measures (Hunter et al., 2023; Mahdi et al., 2018). GVBS is implemented in MATLAB by Harel and colleagues (archived at: https://github.com/Pinoshino/gbvs). We used the saliency map based on intensity (grayscale level), colour, orientation, and motion feature maps. The individual feature maps were combined using equal weights and then normalised to a value between 0 and 1. For the parameter settings, we used a feature map resolution level = 3; the Derrington-Krauskopf-Lennie colour space; four Gabor-filter orientations (0°, 45°, 90° and 135°); and four motion directions (0°, 45°, 90° and 135°). Default settings were used for all other parameters.

For visual clutter, we used the image-based Feature Congestion measure proposed by (Rosenholtz et al., 2007) and implemented in MATLAB (https://dspace.mit.edu/handle/1721.1/37593). This measure computes clutter based on the local variance of several features at multiple scales. These features include colour (CIELab colour space), intensity, and orientation contrasts. Default parameter settings were used.

We down-sampled the video data to 3 fps from 30 fps (i.e., every ten frames). We reduced each frame dimension by 50% (i.e., 320 x 240 pixels) using bicubic interpolation to manage computer memory and computation time. The saliency and clutter map were computed for each frame in the down-sampled and resized video data. The motion feature map for saliency was based on the pixel grayscale difference between every ten frames (∼333 ms).

**Table S2.** Characteristics of binary fixation masks as a function of age. The mean number of fixations used to create the binary fixation mask for each 5-sec time window. The mean percentage of the video frame area taken up by the mask (640 x 480 pixels). The standard deviation (SD) is in brackets.

|  | **Binary Fixation Mask Characteristics** |  |
| --- | --- | --- |
| **Age Group** | **Mean # Fixations (SD)** | **Mean % Frame Area (SD)** |
| 6-months | 6.6 (2.3) | 8.1 (1.8) |
| 9-months | 6.7 (2.6) | 8.1 (2.5) |
| 12-months | 6.5 (2.2) | 7.6 (1.6) |
| 24-months | 6.1 (2.4) | 7.3 (1.9) |
| 36-months | 6.7 (1.8) | 8.4 (2.1) |

**Figure S3.**
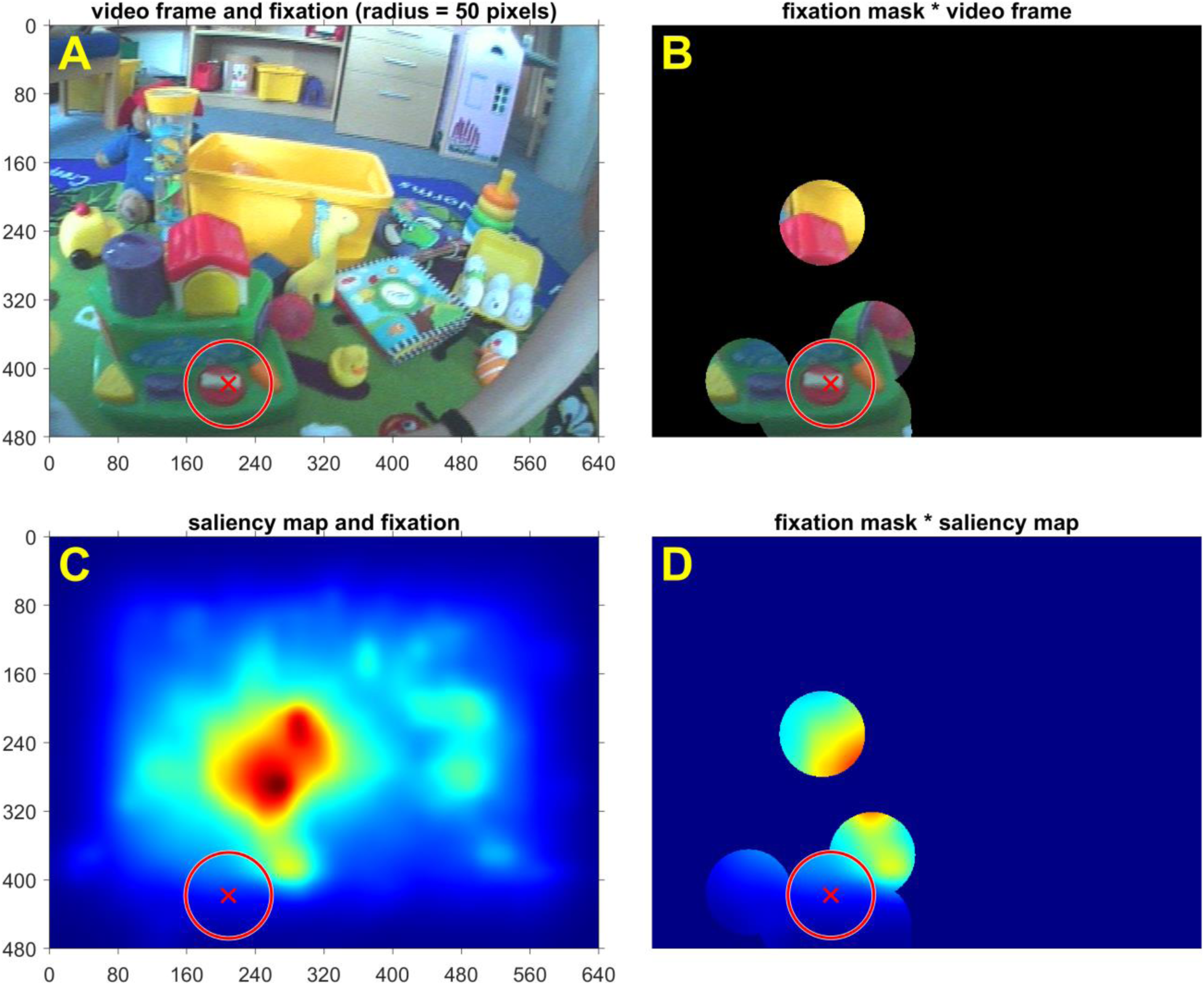
Example of a binary fixation mask applied to a video frame and saliency map. (A) Video frame (time point) with a fixation. The red x is an example fixation, and the red circle represents the radius (*r* = 50 pixels) around that fixation. The same fixation is illustrated in B, C and D. (B) An illustration of applying the binary fixation mask to the video frame. The mask is formed by combining 10 fixations (some fixations overlap each other). (C) The saliency map for the video frame. (D) An illustration of applying the binary fixation mask to the saliency map. The mean saliency for this frame (time point) is the average across all the pixels within the binary fixation mask.

### 6. Change Point Detection

#### The Wild Binary Segmentation2 approach

Typical change point searches amount to the iterative application of algorithmic steps that evaluate a cost function which assumes some data feature knowledge while also embedding a penalty for model complexity (see (Haynes et al., 2017), and references within). Methods pertaining to (wild) binary segmentation have also been proposed and proved to be popular in conventional setups for signals with infrequent changes (Fryzlewicz, 2014). These approaches are designed to detect the most prominent change point and then to reiterate this procedure on the data to its left and right. However, certain data contexts are characterised by the presence of many change points, e.g., economic indicators, such as house price indices; this is also true for our data context here, characterised by the short spans of attention that infants are typically capable of exhibiting. Methodologies driven by common or even adaptive penalties, as well as binary segmentation, have been shown to completely crumble under data scenarios that feature many change points (see (Fryzlewicz, 2020) section 2.1). A way to bypass this issue is to propose a new approach for wild binary segmentation (WBS2) and use a different model selection criterion, namely the steepest drop to low levels methodology (SDLL). This approach is shown to yield excellent results under testing data scenarios pertaining to frequent change point setups (Fryzlewicz, 2020).

The WBS2 has been developed stemming from the WBS (Fryzlewicz, 2014) to specifically address the many change point issues. In order to detect a putative change point in a stretch of data *X*_*s*_, …, *X*_*e*_ (acronyms are for ‘s’=start and ‘e’=end), a binary segmentation (BS) approach computes a CUSUM-type ‘locator’ statistic over the segment [*s*, *e*] and retains the location *s* ≤ *b* < *e* that maximises this statistic.

Mathematically, the true, unknown signal *f*, where *X*_*t*_ = *f*_*t*_ + ɛ_*t*_, is considered to potentially exhibit a change point in *f*_*s*_, …, *f*_*e*_ at *f*_*b*_ for *b* = argmax_*b*_| *CUSUM*_*s,b,e*_(*X*) |, where the CUSUM statistic over the observations in the interval [*s*, *e*] is defined at each location *b* in that interval as

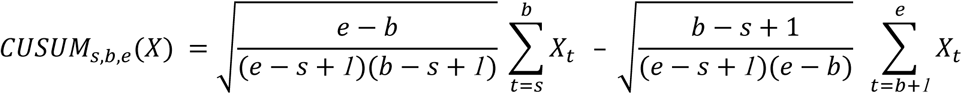

The WBS relies on subsampling the data *X*_1_, …, *X*_T_ over time intervals [*s*, *e*] that are independently, randomly drawn with replacement and giving rise to *M* subsamples *X*_*sm*_, …, *X*_*em*_, *m* = 1, …, *M*. The WBS introduces a further step in which the above maximisation of the CUSUM-locator statistic is computed across the *M* randomly pre-drawn intervals that satisfies [*s*_*m*_, *e*_*m*_] ⊆ [*s*, *e*], hence furthermore *b* = argmax_*sm*,*b,em*_ | *CUSUM*_*sm*,*b,em*_(*X*) |. The identified location is then considered to be a change point if its corresponding CUSUM value exceeds the universal threshold (Donoho & Johnstone, 1994), σ √2 log T, with the noise standard deviation (σ) estimated using the median absolute deviation (MAD) across the entire dataset *X*_1_, …, *X*_T_. Note that the use of thresholding is heavily reliant on a good estimator of σ and when dealing with data that features frequent change-points, this becomes a problem for the performance of WBS. If the putative change-point is deemed significant, then the steps above are re-iterated to its left [*s*, *b*] and right [*b* + 1, *e*] intervals, using the originally drawn subsamples.

When applied to data featuring frequent change points, WBS methodology is required to address (i) the incompleteness of the solution path induced by the use of pre-drawn intervals and (ii) a new model selection strategy. The solution, referred to as WBS2, (i) adaptively uses drawn intervals at each stage based on the location of the detected change-point candidate that maximises the absolute values of the CUSUMs, hence yielding a complete solution path (each time location is a potential candidate in the solution path), and (ii) selects the appropriate change-point using a ‘Steepest-Drop to Low Levels’ criterion (see (Fryzlewicz, 2020) for a complete description).

#### Choice of Changepoint Detection methods

The framework for change point detection consists of modelling the observed (heart rate) data *X*_1_, …, *X*_T_ as a noise-contaminated version of a deterministic piecewise constant function *f* defined over the interval [1, T] whose number of change points (*N*) and their precise locations are unknown. Their reliable estimation is the scope of change point detection procedures across the statistical literature (Truong et al., 2020). Mathematically, this amounts to formalising a model for the observed data *X*_*t*_ = *f*_*t*_ + ε_*t*_, with the noise ε_*t*_ assumed to follow a *N*(0, σ^2^) distribution, which albeit simple still poses major statistical problems, particularly when *N* is large (i.e., there are frequent change points in the true signal *f*) (Fryzlewicz, 2020).

For the work in this paper, 37 separate change point approaches were tested from four separate packages (one from Python, three from R). These were all individually run on a subset of the data (N = 16). The optimal approach was chosen qualitatively by considering the density and positioning of change points across the signal. The heart rate will naturally change in response to a variety of stimuli in addition to attention, and so a method that slightly overpopulates change points was preferred to a more conservative approach. Many approaches use the PELT approach (Pruned Exact Linear Time) (Killick et al., 2012), but the precise choice of algorithm and penalty varies between packages.

From the Ruptures Python package (version 1.1.7) (Truong et al., 2020), the PELT model was used in combination with three penalty types - an L1 model (capturing changes in the median heart rate), an L2 model (capturing changes in the mean heart rate), and a custom model looking at linear changes in slope. All used a log(*L*)σ^2^ penalty value, where *L* is the length of the signal and σ^2^ is the variance.

The Changepoint R package (version 2.2.3) (Killick & Eckley, 2014) uses a PELT algorithm with parametric cost functions. Three approaches were considered from this package, looking at changes in mean, variance, and a combination of mean and variance.

The changepoint.np R package (version 1.0.3) is an extension of the changepoints package that uses the PELT algorithm with nonparametric cost functions based on the empirical distribution of the data but allows for multiple options for penalty. Seven approaches were used, which all considered separate penalties. One approach used no penalties; two approaches used hard penalties (10 and 100), and the other four used penalties criteria-AIC (Akaike information criterion) (Akaike, 1974), SIC (Schwarz information criterion) (Schwarz, 1978), MBIC (modified Bayes information criterion) (Zhang & Siegmund, 2007) and Hannan-Quinn (Hannan & Quinn, 1979) (Package reference - https://www.jstatsoft.org/article/view/v058i03).

The last package considered was the breakfast R package (version 2.2), which uses a two-stage procedure combining a solution path generation with model selection. Four solution paths were combined with six models to produce 24 approaches. The solution paths were: SDLL (steepest descent to lowest levels) (Fryzlewicz, 2020), Strengthened Schwarz information criterion (Baranowski et al., 2019), localised pruning (Cho & Kirch, 2021), and thresholding. The six models were: WBS (Wild Binary Segmentation) (Fryzlewicz, 2014), WBS2 (Fryzlewicz, 2020), iDetect (Anastasiou & Fryzlewicz, 2021), Sequential iDetectSeq (Anastasiou & Fryzlewicz, 2021), Narrowest-Over-Threshold (Baranowski et al., 2019), and TGUH (Tail-Greedy Unbalanced Haar) (Fryzlewicz, 2018) (Package reference https://rdrr.io/cran/breakfast/man/breakfast-package.html).

Out of all of these options, the use of SDLL solution path with WBS2 model in the breakfast R package was chosen due to producing an appropriate density of changepoints for the problem of attention estimation, and also after inspection of these change points by psychologists to confirm that those heart rate changes detected by the approach were of a suitable magnitude.

Other than the nonparametric change point approach with no penalty, almost all other methods were more conservative in change point production than SDLL with WBS2. The choice of SDLL for the solution path was found to be more impactful than the choice of model. The sequential iDetect and TGUH approaches were similar in terms of change point production when combined with the SDLL path, with the WBS2 approach being chosen as the option the package creator recommended for best performance for SDLL. The linear Ruptures approach produced very good localization of change points, but was too sparse for the purposes of attention. Some of the nonparametric change point penalties (MBIC, AIC) were also very close in terms of performance.

### 7. Feature selection using Lasso regularised logistic regression

Feature selection was carried out using Lasso regularised logistic regression, with the following mathematical formulation:

We define a binary response variable *Y* that takes the value 1 if the time point *i* corresponds to attention and 0 otherwise, and model the corresponding probability of attention, denoted by *p*_*i*_, as logit(*p*_*i*_) = *X_i_*^T^β, where 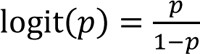 is the (column) vector of covariates corresponding to time *i*, and β is the (column) vector of unknown feature coefficients. Using a suitable transform of the data (denoted here by *Z* and weighted appropriately through a weighted version of the L2 norm, see (Wood, 2017), Section 3.1.2 for a full description), the Lasso approach for the estimation of the regression coefficients yields penalised estimates

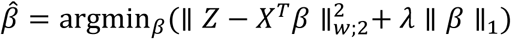

that are exactly zero for those covariates deemed to have insignificant contributions to attention. The estimator sparsity is controlled by the tuning parameter λ, which can be chosen via cross validation.

### 8. Feature selection

Feature selection was conducted using the absolute coefficients from the Lasso regularised logistic regression model. The L1 penalty in Lasso encourages sparsity by setting some coefficients to exactly zero, enabling automatic feature selection. To ensure consistency, we normalised the candidate features and applied 5-fold cross-validation. The mean absolute coefficient values across the folds were used as the criterion for feature selection. An optimal threshold was determined, above which features were selected, with the goal of maximising the F1-score across different threshold values.

**Table S3.** Feature importance and selection criteria. The mean of the absolute coefficients from the Lasso regularised logistic regression model, calculated across the 5-fold cross-validation, was compared to a threshold for feature selection. Selected features are indicated in bold. We also provide the centre frequencies corresponding to the wavelets for WPT-HR-x, WPT-Acc-x, and LSW-HR-Acc-x. WPT-HR: wavelet packet transform of HR; WPT-Acc: wavelet packet transform of Acc; LSW-HR-Acc: local stationary wavelet estimated coherence between HR and Acc.

| # | Feature name | Absolute coefficient* | Notes |
| --- | --- | --- | --- |
| 1 | **HR** | 1.975 |  |
| 2 | Acc | 0.030 |  |
| 3 | **WPT-HR-1** | 1.369 | 0.0312 Hz |
| 4 | **WPT-HR-2** | 0.595 | 0.0938 Hz |
| 5 | **WPT-HR-3** | 0.500 | 0.1562 Hz |
| 6 | **WPT-HR-4** | 0.051 | 0.2188 Hz |
| 7 | **WPT-HR-5** | 0.068 | 0.2812 Hz |
| 8 | **WPT-HR-6** | 0.332 | 0.3438 Hz |
| 9 | **WPT-HR-7** | 0.402 | 0.4062 Hz |
| 10 | **WPT-HR-8** | 0.239 | 0.4688 Hz |
| 11 | **WPT-HR-9** | 0.038 | 0.5312 Hz |
| 12 | **WPT-HR-10** | 0.374 | 0.5938 Hz |
| 13 | WPT-HR-11 | 0.009 | 0.6562 Hz |
| 14 | **WPT-HR-12** | 0.181 | 0.7188 Hz |
| 15 | WPT-HR-13 | 0.016 | 0.7812 Hz |
| 16 | WPT-HR-14 | 0.026 | 0.8438 Hz |
| 17 | **WPT-HR-15** | 0.093 | 0.9062 Hz |
| 18 | **WPT-HR-16** | 0.066 | 0.9688 Hz |
| 19 | WPT-Acc-1 | 0.011 | 0.0312 Hz |
| 20 | WPT-Acc-2 | 0.013 | 0.0938 Hz |
| 21 | WPT-Acc-3 | 0.013 | 0.1562 Hz |
| 22 | WPT-Acc-4 | 0.006 | 0.2188 Hz |
| 23 | WPT-Acc-5 | 0.003 | 0.2812 Hz |
| 24 | WPT-Acc-6 | 0.003 | 0.3438 Hz |
| 25 | WPT-Acc-7 | 0.014 | 0.4062 Hz |
| 26 | WPT-Acc-8 | 0.008 | 0.4688 Hz |
| 27 | WPT-Acc-9 | 0.004 | 0.5312 Hz |
| 28 | WPT-Acc-10 | 0.008 | 0.5938 Hz |
| 29 | WPT-Acc-11 | 0.003 | 0.6562 Hz |
| 30 | WPT-Acc-12 | 0.002 | 0.7188 Hz |
| 31 | WPT-Acc-13 | 0.002 | 0.7812 Hz |
| 32 | WPT-Acc-14 | 0.006 | 0.8438 Hz |
| 33 | WPT-Acc-15 | 0.005 | 0.9062 Hz |
| 34 | WPT-Acc-16 | 0.001 | 0.9688 Hz |
| 35 | LSW-HR-Acc-1 | 0.019 | 0.75 Hz |
| 36 | LSW-HR-Acc-2 | 0.012 | 0.375 Hz |
| 37 | LSW-HR-Acc-3 | 0.023 | 0.1875 Hz |
| 38 | **LSW-HR-Acc-4** | 0.037 | 0.0938 Hz |
| 39 | LSW-HR-Acc-5 | 0.018 | 0.0469 Hz |
| 40 | **LSW-HR-Acc-6** | 0.048 | 0.0234 Hz |
| 41 | LSW-HR-Acc-7 | 0.014 | 0.0117 Hz |
| 42 | LSW-HR-Acc-8 | 0.028 | 0.0059 Hz |
| 43 | **LSW-HR-Acc-9** | 0.051 | 0.0029 Hz |
| 44 | LSW-HR-Acc-10 | 0.018 | 0.0015 Hz |
| 45 | **LSW-HR-Acc-11** | 0.060 | 0.0007 Hz |
| 46 | **LSW-HR-Acc-12** | 0.040 | 0.0004 Hz |
| 47 | **Duration** | 0.465 |  |
| 48 | **Latency** | 0.188 |  |
| 49 | **SDRR** | 0.113 |  |
| 50 | **Age** | 0.120 |  |
| 51 | **Change point binary** | 1.808 |  |

\*Threshold = 0.03

### 9. Alternative classifiers

We evaluated the performance of alternative classifiers trained on the same SMOTE-oversampled training set and tested on a consistent test set. The classifiers examined included logistic regression (used in this study), linear discriminant analysis (LDA), quadratic discriminant analysis (QDA), naive Bayes, K-nearest neighbours (KNN), random forest, and neural network. Overall, their performance was comparable, with only minor differences in the balance between precision and recall.

**Figure S4.**
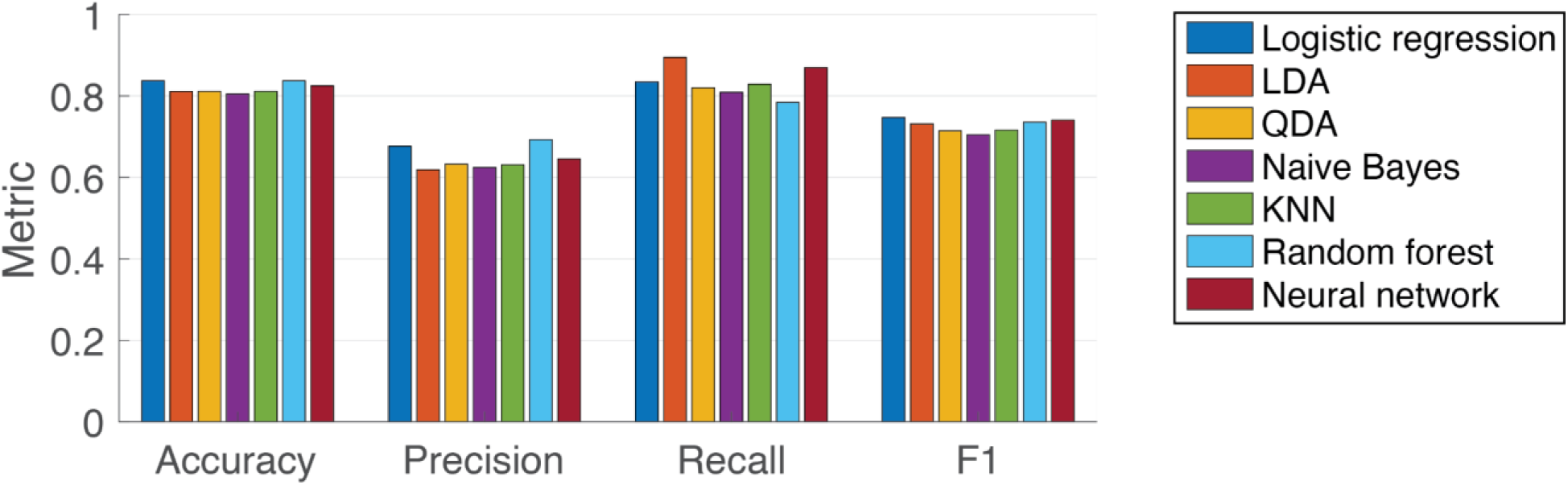
Performance comparison of various classifiers. KNN: 100 neighbours. Neural network: 2 layers, 10 neurons each. Specific values are presented in Table S4.

**Table S4.** Performance metrics of alternative classification models. Metrics were calculated based on the test sets.

| Classifier | Accuracy | Precision | Recall | F1 score |
| --- | --- | --- | --- | --- |
| Logistic regression | 0.8373 | 0.6765 | 0.8345 | 0.7472 |
| LDA | 0.8107 | 0.6187 | 0.8942 | 0.7314 |
| QDA | 0.8108 | 0.6326 | 0.8199 | 0.7142 |
| Naive Bayes | 0.8046 | 0.6243 | 0.8091 | 0.7048 |
| KNN | 0.8108 | 0.6307 | 0.8288 | 0.7163 |
| Random forest | 0.8373 | 0.6924 | 0.7841 | 0.7354 |
| Neural network | 0.8247 | 0.6454 | 0.8694 | 0.7408 |

### 10. Sensitivity test

To evaluate whether the dataset size was adequate for training the model, we conducted a sensitivity test by training on varying fractions of the dataset and assessing performance on the test set. Results showed that precision, recall, and F1 score stabilized when using more than half of the training set, indicating that our model was trained with a sufficient sample size (Figure S5).

**Figure S5.**
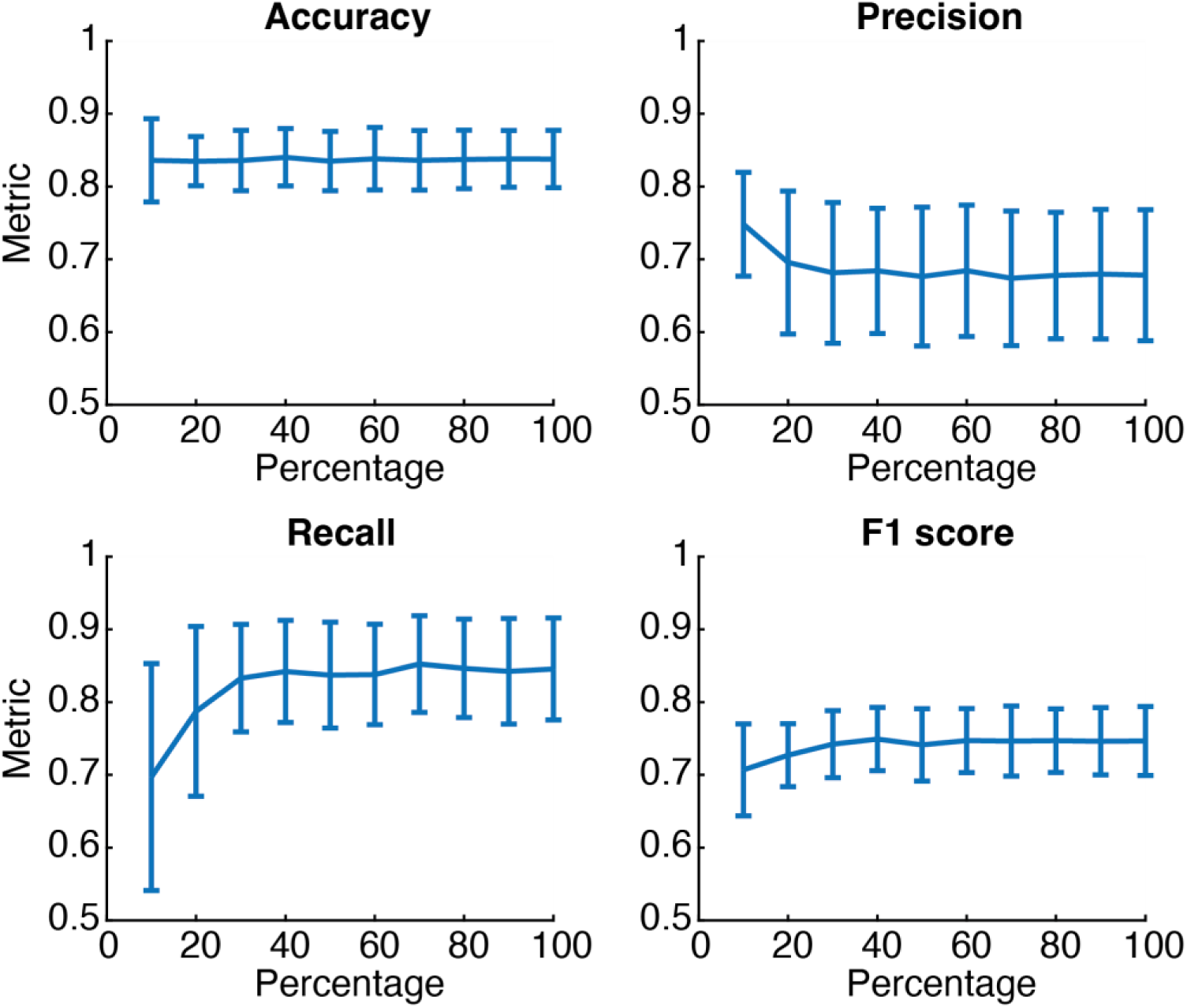
Sensitivity analysis of sample size. The X-axis represents the percentage of the training set used for model training, with evaluations performed on the test set. Lines indicate mean values, and error bars represent standard deviations.

### 11. Statistics of model application

To evaluate ASAP’s effectiveness in detecting natural statistics associated with the attention and inattention states, we extracted the mean visual saliency and mean visual clutter within periods of these two states from each session. We then tested the significance of the following linear model:

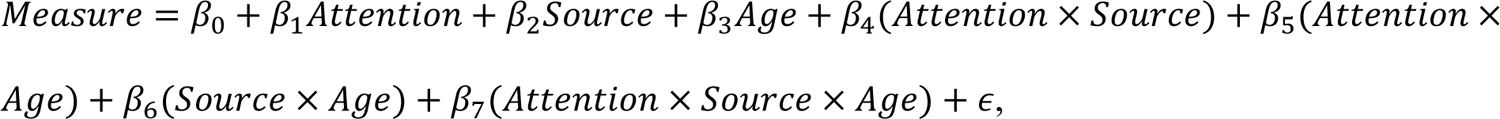

where Measure is either Mean Saliency or Mean Clutter. The p-values were compared to a Bonferroni-corrected significance level α = 0.05/14 = 0.0036, as there were 14 hypotheses (7 coefficients x 2 dependent variables (saliency and clutter)) to test.

**Table S5.**
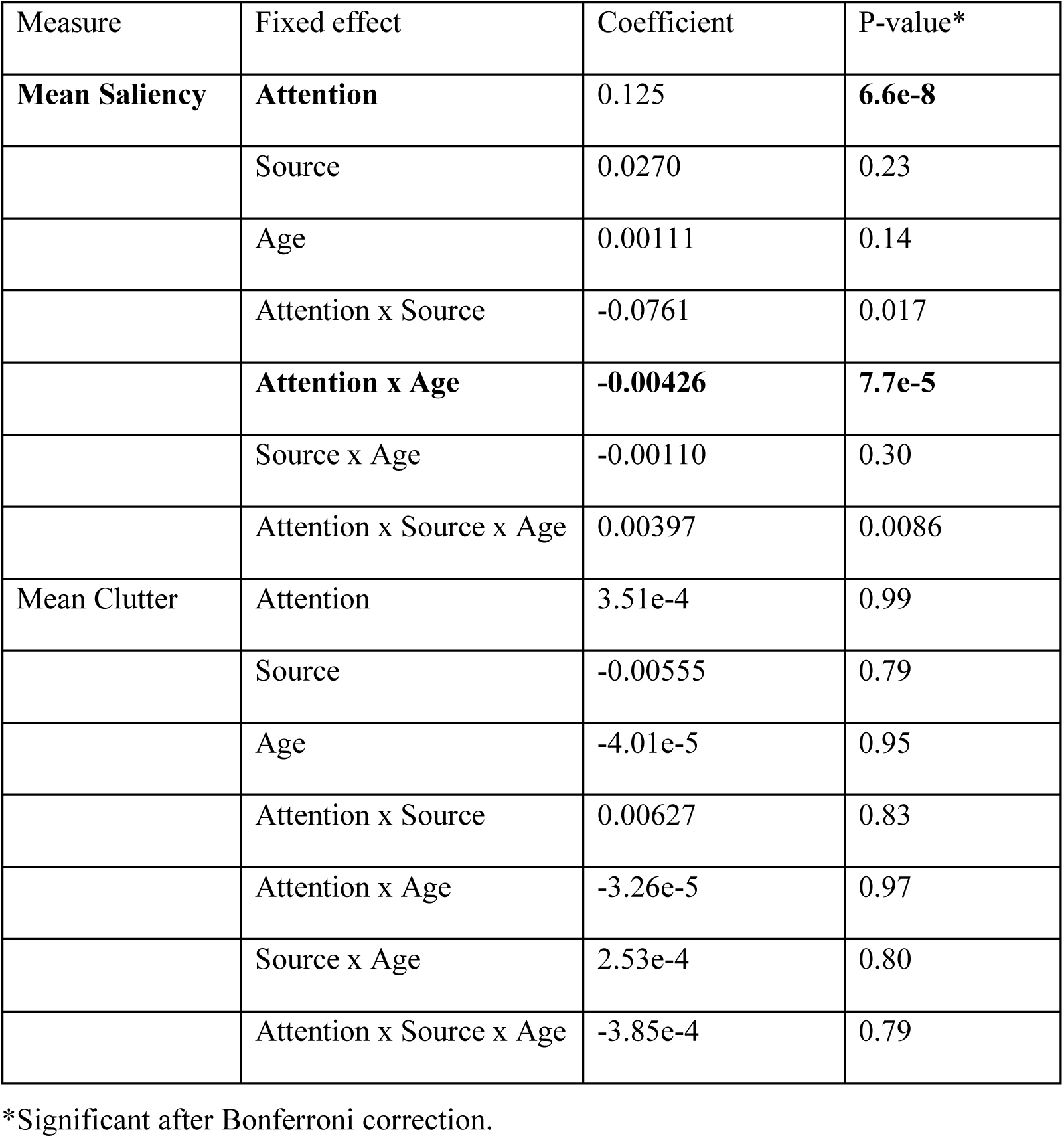
Analysis of variance table for model application. The coefficients and p-values were estimated by fitting a linear model.

| Measure | Fixed effect | Coefficient | P-value* |
| --- | --- | --- | --- |
| **Mean Saliency** | **Attention** | 0.125 | **6.6e-8** |
|  | Source | 0.0270 | 0.23 |
|  | Age | 0.00111 | 0.14 |
|  | Attention x Source | -0.0761 | 0.017 |
|  | **Attention x Age** | **-0.00426** | **7.7e-5** |
|  | Source x Age | -0.00110 | 0.30 |
|  | Attention x Source x Age | 0.00397 | 0.0086 |
| **Mean Clutter** | Attention | 3.51e-4 | 0.99 |
|  | Source | -0.00555 | 0.79 |
|  | Age | -4.01e-5 | 0.95 |
|  | Attention x Source | 0.00627 | 0.83 |
|  | Attention x Age | -3.26e-5 | 0.97 |
|  | Source x Age | 2.53e-4 | 0.80 |
|  | Attention x Source x Age | -3.85e-4 | 0.79 |

\*Significant after Bonferroni correction.

### 12. Source of discrepancies between human coder and ASAP labels

The observed variability in performance across sessions prompted us to investigate whether these fluctuations were attributable to the intrinsic characteristics of the dataset or the model’s inherent biases. In the first scenario, data with specific characteristics would consistently yield better performance, regardless of whether the labels are from humans or machines. In the second scenario, the model might perform well with certain data types, even though human coders show different consistency.

To test this, we examined whether the level of agreement between human coders (HH) correlated with the agreement between a human coder and the ASAP model (HM). If data characteristics drove the performance variation, we would expect a correlation between HH and HM agreements. For HH agreement, we calculated Cohen’s κ between two human coders (see *Inter-rater reliability of human coding*). We calculated Cohen’s κ using the same participant sessions (N = 15 subjects) as the HH calculation for HM agreement. The HM agreement was assessed between the machine’s predicted attention and Human Coder #1’s annotations, which were used for model training.

Our initial correlation analysis revealed a statistically significant positive correlation between HH and HM Cohen’s κ values (Pearson’s *r* = 0.55, *p* = 0.017, permutation test; Figure S5A, B), suggesting that specific dataset characteristics might influence performance variability. To investigate potential underlying factors driving this correlation, we examined age, heart rate variability (assessed via SDRR), change point density (number of change points detected per second), and the extent of movement (assessed via the standard deviation of Acc). We tested whether the correlation between HH and HM Cohen’s κ was attributable to these factors by calculating partial correlations conditioned on each. Except for SDRR (*p* = 0.074, permutation test), the partial correlations with other factors remained significantly greater than 0 (*p* < 0.05; Figure S5C). These results suggest that HRV, as measured by SDRR, influenced the uncertainty of attention detection for both humans and ASAP. Specifically, higher HRV was associated with increased inter-rater reliability in identifying periods of attention (Pearson’s correlations *r* = 0.42 between SDRR and HH, and *r* = 0.55 between SDRR and HM Cohen’s κ). Other possible sources of discrepancies are discussed in the Discussion section but are beyond the scope of the current study.

**Figure S6.**
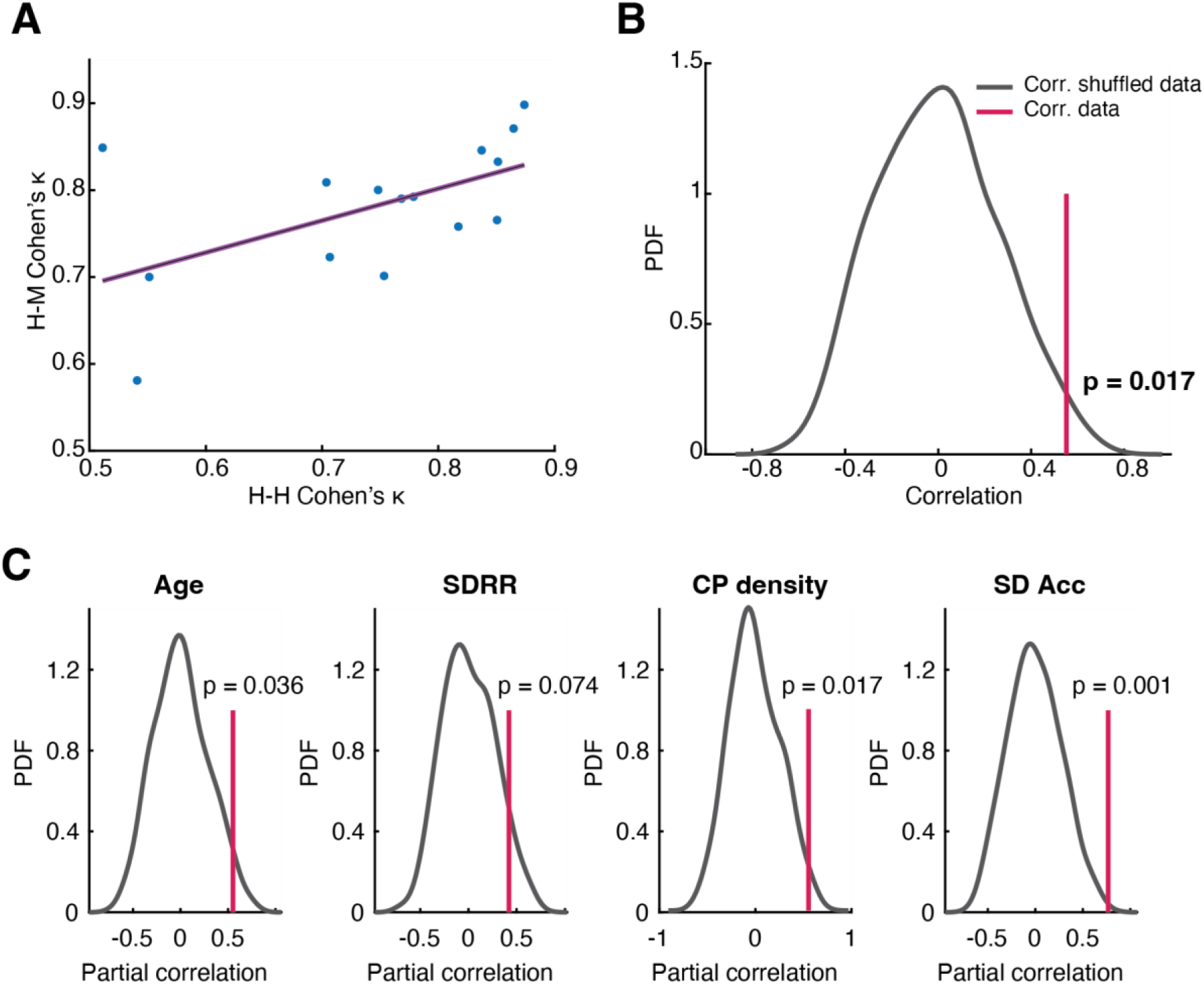
Analysis of the source of disagreement between human coders and ASAP. (A) Scatter plot of H-M vs. H-H Cohen’s κ. The line is the linear fit. (B) Distribution of the HH-HM correlation with shuffled data. The red line indicates the data’s correlation. (C) Shuffled data establish the null distributions of partial correlations. Red lines indicate the data’s partial correlation conditioned on each factor. PDF: probability density function. CP: change point.

## Notes

### Competing Interest Statement

The authors have declared no competing interest.

